# The Genomic Legacy of Aurochs hybridization in ancient and modern Iberian Cattle

**DOI:** 10.1101/2023.07.19.548914

**Authors:** Torsten Günther, Jacob Chisausky, M. Ángeles Galindo-Pellicena, Eneko Iriarte, Oscar Cortes Gardyn, Paulina G. Eusebi, Rebeca García-González, Irene Urena, Marta Moreno, Alfonso Alday, Manuel Rojo, Amalia Pérez, Cristina Tejedor Rodríguez, Iñigo García Martínez de Lagrán, Juan Luis Arsuaga, José-Miguel Carretero, Anders Götherström, Colin Smith, Cristina Valdiosera

## Abstract

Cattle have been a valuable economic resource and cultural icon since prehistory. From the initial expansion of domestic cattle into Europe during the Neolithic period, taurine cattle (*Bos taurus*) and their wild ancestor, the aurochs (*B. primigenius*), had overlapping ranges, leading to ample opportunities for mating (whether intended by farmers or not). We performed a bioarchaeological analysis of 24 Bos remains from Iberia dating from the Mesolithic to the Roman period. The archaeogenomic dataset allows us to investigate the extent of domestic-wild hybridization over time, providing insight into the species’ behavior and human hunting and management practices by aligning changes with cultural and genomic transitions in the archaeological record. Our results show frequent hybridization during the Neolithic and Chalcolithic, likely reflecting a mix of hunting and herding or relatively unmanaged herds, with mostly male aurochs and female domestic cattle involved in hybridization. This is supported by isotopic evidence consistent with ecological niche sharing, with only a few domestic cattle possibly being managed. The proportion of aurochs ancestry remains relatively constant from about 4000 years ago, probably due to herd management and selection against first generation hybrids, coinciding with other cultural transitions. The constant level of wild ancestry (∽20%) continues into modern western European breeds including the Spanish Lidia breed which is bred for its aggressiveness and fighting ability, but does not display elevated levels of aurochs ancestry. This study takes a genomic glance at the impact of human actions and wild introgression in the establishment of cattle as one of the most important domestic species today.

## Introduction

Domestication of livestock and crops has been the dominant and most enduring innovation of the transition from a hunter-gathering lifestyle to farming societies. It represents the direct exploitation of genetic diversity of wild plants and animals for human benefit. Ancient DNA (aDNA) has proved crucial to understanding the domestication process and the interaction between domesticated species and their wild relatives both within domestication centers and throughout the regions that the domestics expanded into (*1–14*). The origins of the European domestic taurine, *Bos taurus*, are located in the Fertile Crescent (*15, 16*) and unlike dogs, pigs and goats, where the wild forms are still extant, the wild cow (the aurochs) went extinct in 1627. Aurochs, *B. primigenius*, was present throughout much of Eurasia and Africa before the expansion of domestic cattle from the Levant that accompanied the first farmers during the Neolithisation of Europe. Upon arrival, these early incoming domesticates inevitably coexisted with their wild counterparts in great parts of Europe facilitating gene flow in both directions. In general, taxa within the genus *Bos* can hybridize and produce fertile offspring (*17*) which may have facilitated and contributed to domestication, local adaptation and even speciation (*5, 18–20*). Mitochondrial DNA studies have previously indicated gene flow between domestic cattle and aurochs outside their domestication center (*21–25*) and more recently, genomic studies have shown the presence of European aurochs ancestry in modern taurine cattle breeds (*26–28*). Although cattle have represented a significant economic resource and a prominent cultural icon for millennia, and have been studied through aDNA for more than a decade (*5, 21, 22, 26, 28–30*), our understanding of the interaction of early cattle herds and wild aurochs is still limited due to a lack of time-series genomic data. This gap of knowledge includes European aurochs’ genetic contribution to modern domestic breeds and human management of these animals in the past.

Aurochs have been widely exploited by humans since the European Palaeolithic and archaeological evidence indicates that the species survived in Europe until historical times. Iberia could have served as a glacial refugium for aurochs (*28*), and the most recent evidence for aurochs is found at a Roman site in the Basque Country (*31*). Domestic cattle were introduced into Iberia with the Mediterranean Neolithic expansion and reached the northern coast of the peninsula around 7000 years cal BP (*32*). Consequently, aurochs and domestic cattle have coexisted in Iberia for about five millennia. Since then, cattle have played an important role in Iberian societies as a source of food and labour, as well as cultural events such as bullfighting. Currently, there are more than 50 bovine breeds officially recognized in the Iberian Peninsula including the Lidia breed, a primitive, isolated population selected for centuries to develop agonistic-aggressive responses with the exclusive purpose of taking part in such socio-cultural events (*33*). Recently, it has been reported that Lidia breed individuals have the largest brain size among a comprehensive data set of European domestic cattle breeds and are the most similar to wild aurochs (*34*). The combination of aggressiveness and larger brain size in the Lidia breed may suggest a higher proportion of aurochs ancestry compared to other cattle breeds.

Here, we present the genomes and stable isotope data of Iberian Bovine specimens ranging from the Mesolithic into Roman times from four archeological sites (Fig. 1A). We explore the extent of interbreeding between wild aurochs and domestic cattle over time and the correlation of genetic ancestry with metric identification and ecological niches. Finally, we compare the results to genomic data obtained from modern Iberian cattle breeds to estimate the genetic contribution of the now-extinct aurochs to the Iberian farming economy.

**Figure 1:**
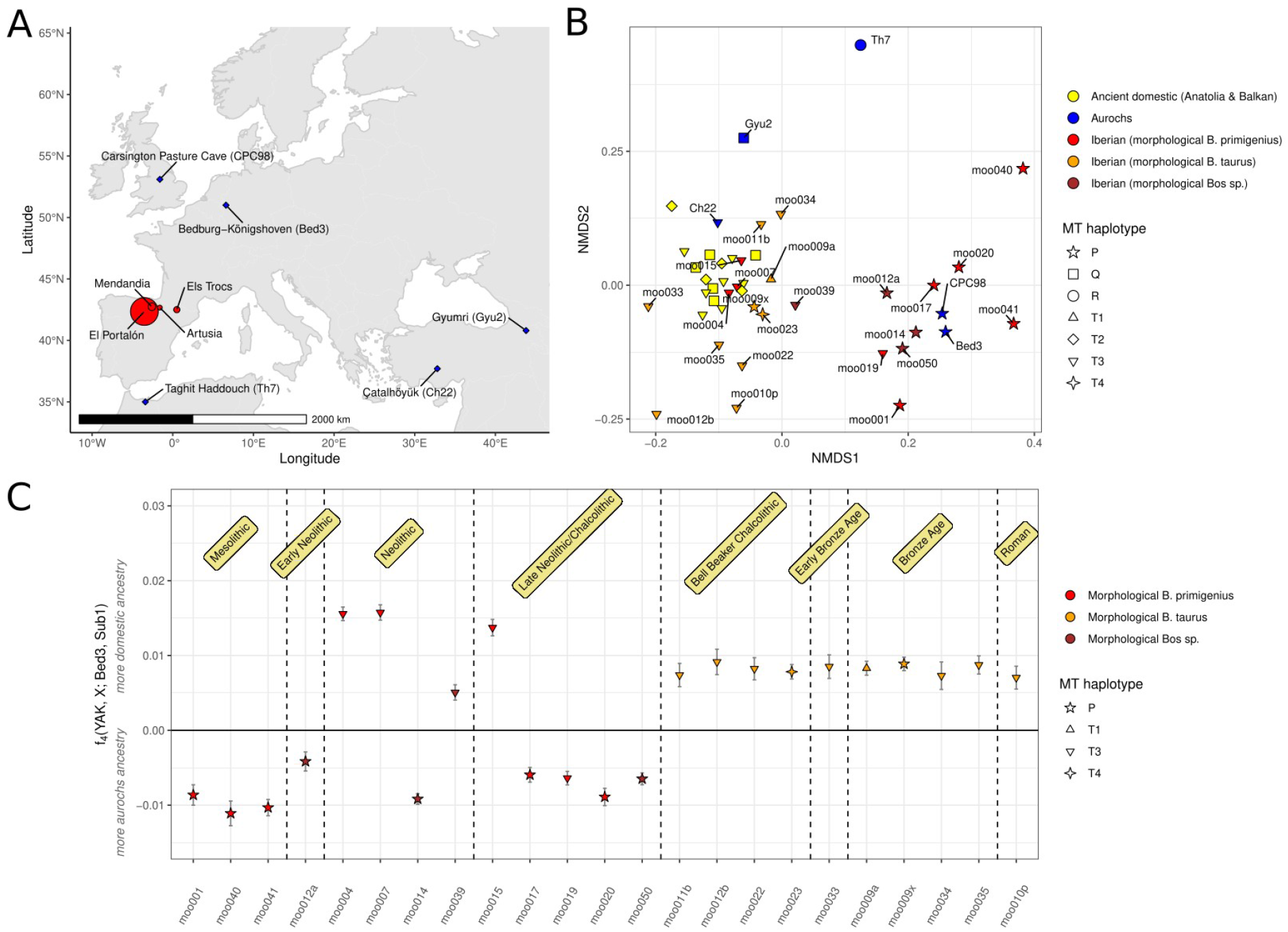
(A) Map of Europe showing the Iberian sampling sites (red circles, size proportional to sample sizes) and the sites for published aurochs genomes used in the analysis (blue diamonds). (B) NMDS ordination of nuclear data of Iberian samples considered *B. taurus* samples (orange), Iberian samples that were morphologically considered *B. primigenius* (red), other Iberian *Bos* samples (brown), ancient domestic cattle from the Balkans and Anatolia (yellow), and aurochs (blue). Data point shape corresponds to mitochondrial haplogroups. (C) f_4_ statistic measuring allele sharing of the Iberian samples with European aurochs (Bed3) or Anatolian Neolithic cattle (Sub1). The time periods displayed are contextual.

## Results

### Exploratory analysis

We successfully sequenced 24 Bovine specimens excavated at four prehistoric sites in Iberia (Fig. 1A). Nine of these individuals were inferred to or suspected to represent aurochs based on morphology or chronology. Direct radiocarbon dates and contextual dating placed the individuals between the Mesolithic (oldest sample moo001, 8641-8450 cal BP) and the Roman Age (youngest sample moo010p, 2260-2150 cal BP). It should be noted that while all post-Mesolithic samples were found at archaeological sites with evidence for herding of other domestic fauna such as ovicaprids (Supplementary Information), we do not know whether these bovids were herded or hunted. Based on the number of reads mapping to the X chromosome, 13 individuals were identified as female and 11 as male, for the samples with sufficient amounts of reads for this analysis. Sequencing coverage of the nuclear cattle genome was low to medium, reaching up to 4.7x with a mean of 0.38x (Dataset S1). The sequence data for non-UDG-treated libraries showed damage patterns characteristic of ancient DNA (Supplementary Information, Figure S1). Based on reads mapping to the mitochondrial genome, we were able to estimate contamination for 11 samples with most samples showing low levels (<2%) of contamination. One individual (moo013a) showed a high contamination of 51.7% [33.5, 69.9] and was excluded from further analysis. One Mesolithic individual (moo040) showed 8.3% [3.1, 13.5] contamination, which was included in the initial exploratory analysis but not used for the analysis of hybridization between wild and domestic as this analysis only focused on post-Mesolithic individuals.

Nine individuals were assigned to the mitochondrial P1 haplogroup, one to haplogroup T1, one to haplogroup T4, and 12 to haplogroup T3 (Dataset S1). P haplogroups are dominant among and thought to be endemic to European aurochs (*35*), but are occurring at low frequencies in modern European cattle breeds (*24*). The prevalence of the T3 haplogroup in our samples is expected; this haplogroup is dominant among modern European *B. taurus* and is the most common haplogroup in ancient Western European domestic cattle. T3 was found in directly dated Neolithic samples from different sites providing direct evidence for the arrival of domestic cattle in northern Iberia during the Neolithic (Dataset S1). The specimen assigned to T1 (moo009a) is notable since this individual was previously used to argue for Bronze Age contact between Iberia and Africa, where the T1 haplogroup is thought to have originated (*29, 36*). T4 is usually considered to be restricted to Asian breeds with rare finds in Europe, restricted to the Balkans (*24*). The presence of T4 in Chalcolithic Iberia suggests that this haplogroup must have been distributed across Western Europe at low frequencies in prehistory. Furthermore, the fact that some specimens that were morphologically identified as aurochs carry domestic T haplogroups implies some level of interbreeding between the two groups.

As mitochondrial genomes only reflect the maternal line of ancestry, they are not informative about the exact extent of interbreeding in our dataset. To avoid being constrained by the variation in modern domestic breeds as with common approaches such as projected PCA (Supplementary Information, Figure S4), we performed non-metric multi-dimensional scaling (NMDS) ordinations on a matrix of pairwise outgroup f_3_ statistics to explore the genomic ancestry of the sequenced individuals. For reference, we included early cattle genomes from Anatolia and the Balkans as well as aurochs excavated from Morocco (Th7), Armenia (Gyu2), Anatolia (Ch22), Germany (Bed3) and Britain (CPC98) (*5, 26, 30*), and calculated pairwise outgroup f_3_ statistics. The NMDS ordination outcome (Fig. 1B) seems to represent a separation between domestic autosomal ancestry (to the left) and European aurochs ancestry (to the right). In contrast, aurochs from other regions (Th7 and Gyu2) seem genetically distinct. Many early domestic samples from Iberia fall close to early cattle from the Balkans and Anatolia as well as the Anatolian aurochs (Ch22). Notably, at least two of the Iberian samples in this cluster (moo004, moo007) were morphologically identified as aurochs. Eight of the nine Iberian samples with haplogroup P fall to the right in the plot, together with the aurochs from Germany (Bed3) and Britain (CPC98). Additionally, one individual carrying a domestic T3 mitochondrial genome (moo019) appears closer to the aurochs samples than the domestics. Out of nine samples that were presumed aurochs based on their morphological features, only six would be considered aurochs based on this analysis. This highlights a substantial overlap between measurements or criteria that are used to distinguish wild and domestic *Bos* based on morphometrics.

This analysis suggests that one can use other European aurochs such as the German Bed3 (Bedburg-Königshoven, 11802-11326 CalBP) (*30*) or the British CPC98 (Carsington Pasture Cave, 6874-6602 CalBP) (*26*) as a reference for Western European aurochs as they seem similar to our three low-coverage Mesolithic Iberian samples. This is also supported by a recent parallel study concluding that all Western European aurochs form a clade, possibly even originating from an Iberian glacial refugium (*28*). Using Sub1 (Suberde Höyük, 8171-7974 CalBP) (*5*), a Neolithic domestic Anatolian individual, and the higher coverage aurochs Bed3 as references, we can perform f_4_ statistics to measure which Iberian individuals share more alleles with one or the other (Fig 1C). Despite the relatively low coverage of some samples, the f_4_ statistics are highly correlated with the first axis of the NMDS (R^2^=0.84, p=8.3×10^−10^) implying that they detect the same pattern. Non-overlapping confidence intervals also confirm that the high genetic differentiation between Western European aurochs and domestic cattle allows confident assignment even with low coverage data. The three Mesolithic individuals as well as an additional six, up until the Late Neolithic/Chalcolithic, share most of their alleles with aurochs. Three individuals from the Neolithic and Late Neolithic/Chalcolithic share most of their alleles with domestic Anatolian cattle while two individuals (moo012a and moo039) are more intermediate, suggesting that there could have been some level of hybridization. More recent samples from the Bell Beaker period onwards all appear to have similar amounts of allele sharing with mostly domestic ancestry but some level of aurochs introgression.

### Quantifying the extent of introgression

While f_4_ statistics measure allele sharing it does not directly quantify the amount of introgression in the different specimens, hence, we employed three different frameworks to estimate ancestry proportions: f_4_ ratio (*37*), qpAdm (*38, 39*) and Struct-f4 (*40*) to model each Iberian individual from European aurochs (Bed3) and/or Anatolian Neolithic cattle (Sub1) as sources (Table 1). While the f_4_ ratio provides a straightforward-to-interpret estimate of aurochs ancestry under a simple two source model, we also include qpAdm due to the potential of rejecting models and hinting at additional ancestries. We also include Struct-f4 for better samples (>0.1X) as it is more flexible than qpAdm not requiring a strict separation between sources and outgroup populations. While quantitative estimates of European aurochs ancestry for the 20 post-Mesolithic individuals are somewhat correlated between f_4_ ratio and qpAdm (Spearman’s correlation coefficient rho=0.57, p=0.01), they differ for certain individuals. This highlights differences between the methods, their assumptions about the relationships of sources and outgroups, and their sensitivity to low coverage data. For most parts of this study, we decide to present the f_4_ ratio results but it is important to highlight that our interpretations are based on the general pattern and not on the ancestry estimates for single individuals.

**Table 1:** European aurochs ancestry proportions in post-Mesolithic Iberian *Bos* samples. Square brackets are showing Block-jackknife estimates of the 95% confidence interval. f4 ratio and qpAdm are using Bed3 as source of European aurochs ancestry unless noted otherwise. Footnotes are added when deviations from the two source model were needed. Struct-f4 was run in semi-supervised mode to estimate ancestry in the Iberian samples with K=5 as the different ancestries separated at this point. Only individuals with at least 0.1x coverage were included in this analysis to ensure convergence. LNCA=Late Neolithic/Chalcolithic

| Sample ID | Site | Period | Date calBP | f<sub>4</sub> ratio | qpAdm | Struct-f4 (K=5) |
| --- | --- | --- | --- | --- | --- | --- |
| moo012a | El Portalón | Early Neolithic | *contextual* | 0.908 [0.743, 1.072] | 0.928 [0.77, 1.085] | - |
| moo004 | Els Trocs | Neolithic | 7152-6890 | 0.103 [0.025, 0.182] | 0.06 [0.026, 0.098]<sup>1</sup> | 0.012<sup>a</sup> |
| moo007 | Els Trocs | Neolithic | 7151-6890 | -0.089 [-0.204, 0.026] | 0.073 [0.033, 0.113]<sup>1</sup> | 0.039<sup>a</sup> |
| moo014 | El Portalón | Neolithic | 6491-6403 | 0.861 [0.791, 0.931] | 0.942 [0.905, 0.979]<sup>2</sup> | 0.942<sup>a</sup> |
| moo039 | Mendandía | Neolithic | 7426-7280 | 0.435 [0.334, 0.536] | 0.866 [0.76, 0.971] | - |
| moo015 | El Portalón | LNCA | 5041-4842 | -0.055 [-0.182, 0.072] | 0.444 [0.32, 0.565] | - |
| moo017 | El Portalón | LNCA | 5567-5326 | 0.756 [0.646, 0.866] | 0.811 [0.769, 0.853]<sup>1</sup> | - |
| moo019 | El Portalón | LNCA | 5556-5325 | 0.723 [0.628, 0.818] | 0.828 [0.786, 0.87]<sup>1</sup> | 0.654 |
| moo020 | El Portalón | LNCA | *contextual* | 0.775 [0.652, 0.897] | 0.944 [0.80, 1.085] | - |
| moo050 | El Portalón | LNCA | 5468-5320 | 0.788 [0.709, 0.866] | 0.809 [0.773, 0.845]<sup>1</sup> | 0.663<sup>b</sup> |
| moo011b | El Portalón | Bell Beaker | *contextual* | 0.109 [-0.097, 0.316] | 0.667 [0.47, 0.865] | - |
| moo012b | El Portalón | Bell Beaker | 4421-4291 | 0.706 [0.528, 0.883] | 0.326 [0.258, 0.394]<sup>1</sup> | - |
| moo022 | El Portalón | Bell Beaker | *contextual* | 0.169 [0.006, 0.332] | 0.729 [0.55, 0.910] | - |
| moo023 | El Portalón | Bell Beaker | 4153-3976 | 0.179 [0.073, 0.285] | 0.641 [0.54, 0.738] | 0.21<sup>a</sup> |
| moo033 | El Portalón | Early Bronze Age | *contextual* | 0.241 [0.050, 0.432] | 0.987 [0.79, 1.182] | - |
| moo009a | El Portalón | Bronze Age | 3884-3635 | 0.399 [0.297, 0.502] | 0.284 [0.244, 0.324]<sup>1</sup> | 0.152<sup>a</sup> |
| moo009x | El Portalón | Bronze Age | 3829-3513 | 0.198 [0.115, 0.280] | 0.259 [0.203, 0.315]<sup>3</sup> | 0.176<sup>a</sup> |
| moo034 | El Portalón | Bronze Age | 3811-3492 | 0.547 [0.300, 0.794] | 0.112 [-0.061, 0.285]<sup>4</sup> | - |
| moo035 | El Portalón | Bronze Age | *contextual* | 0.307 [0.170, 0.445] | 0.821 [0.68, 0.963] | - |
| moo010p | El Portalón | Roman | 2334-2156 | 0.397 [0.205, 0.589] | 0.281 [0.217, 0.345]<sup>1</sup> | - |

<sup>1</sup> To produce a fitting and feasible model (p>0.01) a minor contribution (<=5%) of indicine ancestry is required to fit the data.
<sup>2</sup> Model does not fit with Bed3 as European aurochs source but fits well (p=0.49) when using CPC98.
<sup>3</sup> To produce a fitting and feasible model (p>0.01) a contribution of 33.7% Caucasus aurochs ancestry (Gyu2) is required to fit the data. This additional source is not well resolved as the standard error is large (21.2%).
<sup>4</sup> To produce a fitting and feasible model (p>0.01) a contribution of 85.9% Caucasus aurochs ancestry (Gyu2) is required to fit the data. This additional source is not well resolved as the standard error is large (74.6%).
<sup>a</sup> Struct-f4 also assigns a small proportion of Caucasus aurochs ancestry (Gyu2) to this individual (<5%).
<sup>b</sup> Struct-f4 also assigns a proportion of Caucasus aurochs ancestry (Gyu2) to this individual (14.0%).

Most of the 20 post-Mesolithic individuals show indications of both domestic and European aurochs ancestries (Table 1). Only three individuals (f_4_ ratio) or one individual (qpAdm) do not show significant proportions of aurochs ancestry while only one individual (f_4_ ratio) or three individuals show not significant proportions of domestic ancestry. Furthermore, qpAdm and Struct-f4 suggest low proportions of additional, eastern ancestries represented either by indicine cattle or the Caucasus aurochs Gyu2 in these analyses. While these ancestries are not well resolved and usually have high standard errors, they suggest that multiple western Asian populations contributed to the European early domestic gene pool. Notably, most Neolithic and pre-Bell Beaker Chalcolithic individuals show either predominantly domestic or aurochs ancestry while many Bell Beaker and Bronze Age individuals show more intermediate values of aurochs ancestry. In fact, from the Bronze Age onwards, most estimates overlap with the approximately 25% aurochs ancestry in modern Iberian cattle (Fig. 2; Supplementary Information, Table S1) (*41*).

**Figure 2:**
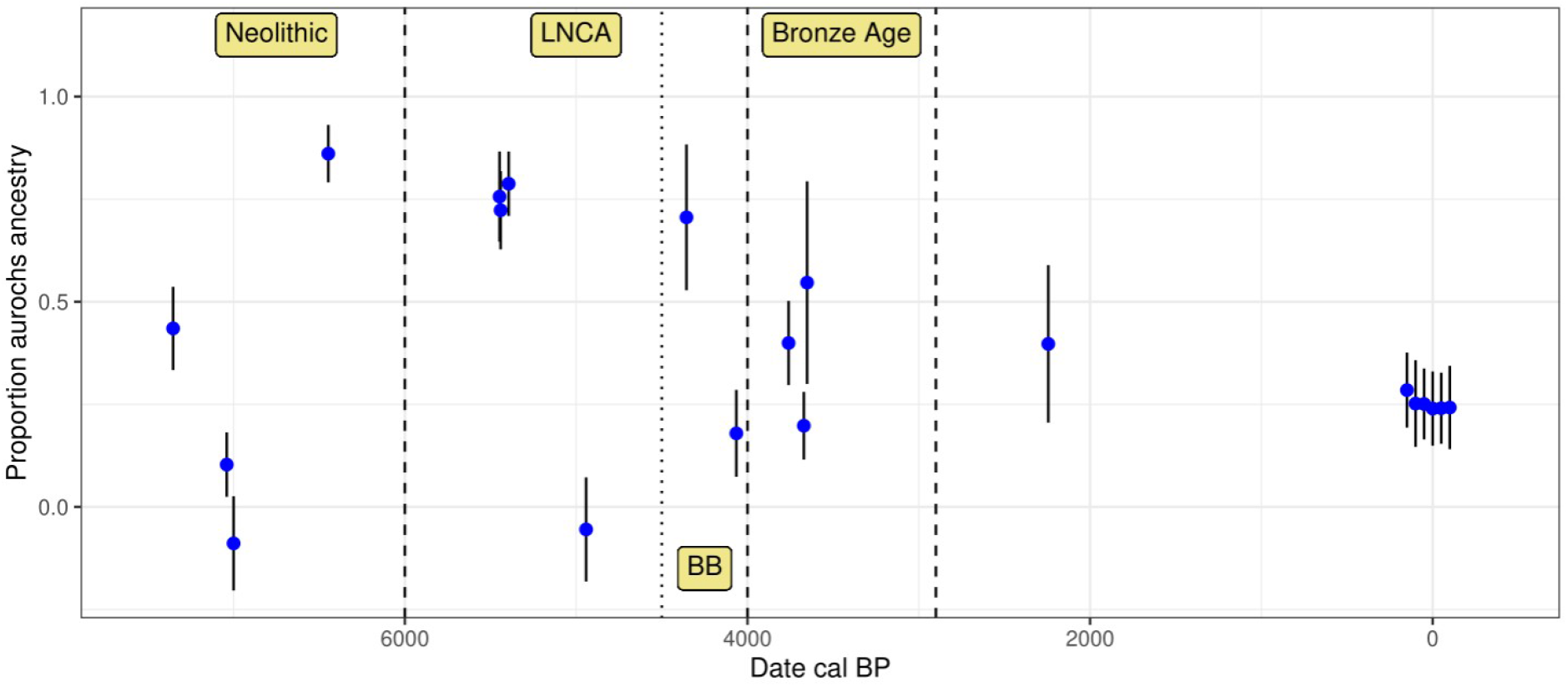
(A) Estimates of aurochs ancestry (estimated using the f_4_ ratio with Bed3 as European aurochs source) in directly dated post-Mesolithic Iberian samples over time. Error bars indicate the 95% confidence interval. Modern Iberian whole-genome sequenced Lidia individuals are added around date 0 with some horizontal jitter. Approximate boundaries for the main sampling periods are indicated by dashed vertical lines.

A limitation of this analysis is the availability of genomes that can be used as representatives of the source populations. We used German and British aurochs to represent western European aurochs ancestry and a single Anatolian Neolithic to represent the original domestic cattle that was introduced into Europe. Our Mesolithic Iberian aurochs contained too little endogenous DNA to be used as a proxy aurochs reference and all Neolithic and Chalcolithic samples estimated with predominantly aurochs ancestry (including the 2.7x genome of moo014) already carry low (but significant) levels of domestic ancestry. However, the fact that all of these aurochs samples carried P mitochondria strongly suggests that western European aurochs can be considered monophyletic. Furthermore, a recent parallel study also concluded that all Western European aurochs form a clade (*28*). The Anatolian Sub1 might also not be depleted of any European aurochs ancestry and could not fully represent the original European Neolithic gene pool as also indicated by qpAdm and Struct-f4 identifying small proportions of other Asian ancestries in some Iberian individuals. While these caveats should affect our quantitative estimates of European aurochs ancestry, they should not drive the qualitative pattern as our tests would still detect any excess European aurochs ancestry that was not present in Neolithic Anatolia.

An important question that remains unexamined is the exact process that led to the hybridization since this could provide insight into human management practices or, more generally speaking, mating patterns between wild and domestic individuals. The fact that some individuals with predominantly aurochs ancestry carry T haplogroups (moo019) and that some individuals with predominantly domestic ancestry carry P haplogroups (moo009x) implies that females contributed in both directions. To assess whether the admixture process was sex-biased, we compared aurochs ancestry patterns on the X chromosome and autosomes (Fig. 3). Since females carry two X chromosomes and males only have one, we can assume that an excess of a certain ancestry on the X chromosome indicates more females from that particular source population. While the estimates are noisy due to the low coverage data and even less sites available for the X chromosome, it is striking that all but one individual with mostly domestic autosomal ancestry (>50%) show even lower point estimates of aurochs ancestry on the X chromosome. This pattern even extends into the modern Iberian individuals. Male-biased aurochs introgression has been suggested based on mitochondrial haplotypes before (*5*). In the absence of aurochs Y chromosomal data, however, it is difficult to assess sex-biased processes from uniparental data alone. The comparison of X chromosomes and autosomes should theoretically have more power to detect such processes as they are less sensitive to genetic drift due to their recombining nature (*42*) but estimation of ancestry proportions on the X chromosome can be affected by different biases (*43–45*). Overall, our results are consistent with previous observations that the contribution of wild ancestry into domestic cattle was mostly through aurochs bulls.

**Figure 3:**
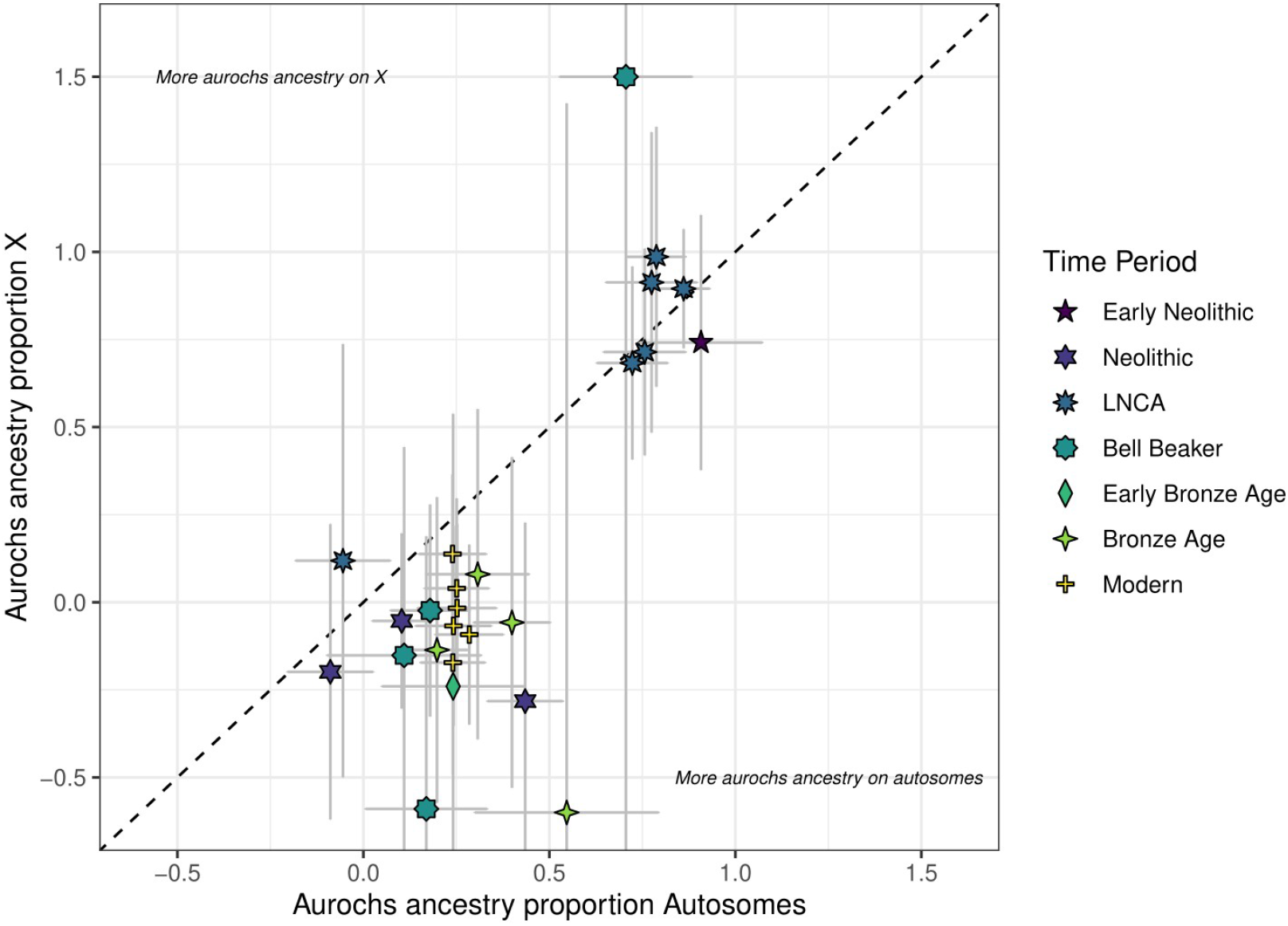
Comparison of f_4_ ratio estimated aurochs ancestry of post-Mesolithic Iberian samples on the autosomes versus X chromosomes. Error bars indicate the 95% confidence interval.

### Aurochs ancestry in modern breeds and the Spanish Lidia cattle breed

We estimated aurochs ancestry in a set of Western European cattle breeds (*27*) as we performed for the prehistoric samples. Previous studies have used *D* statistics for pairwise comparisons between breeds (*26, 27, 46*). Such *D* statistics, however, are sensitive to biases including gene flow from populations not included in the analysis (*47*). Furthermore, qpAdm provides the possibility to reject scenarios not fitting the data. Our point estimates for the aurochs ancestry range between 20% and 30% across all breeds (Fig. 4) and do not show an increase in aurochs ancestry in Iberian breeds (*46*). This result differs from the previous studies which suggested geographic differences in western and central Europe and we believe this could be due to ancestry from other, non-European groups in some commercial breeds (Supplementary Information). Importantly, not all tested breeds did fit the simple two-source model Anatolian Neolithic domestic + European aurochs, likely representing low levels of contributions from other groups, e.g. indicine cattle (*27*). The presence of indicine ancestry can be confirmed in a qpAdm analysis using three sources resulting in fitting models for all breeds (Supplementary Information, Table S4).

**Figure 4:**
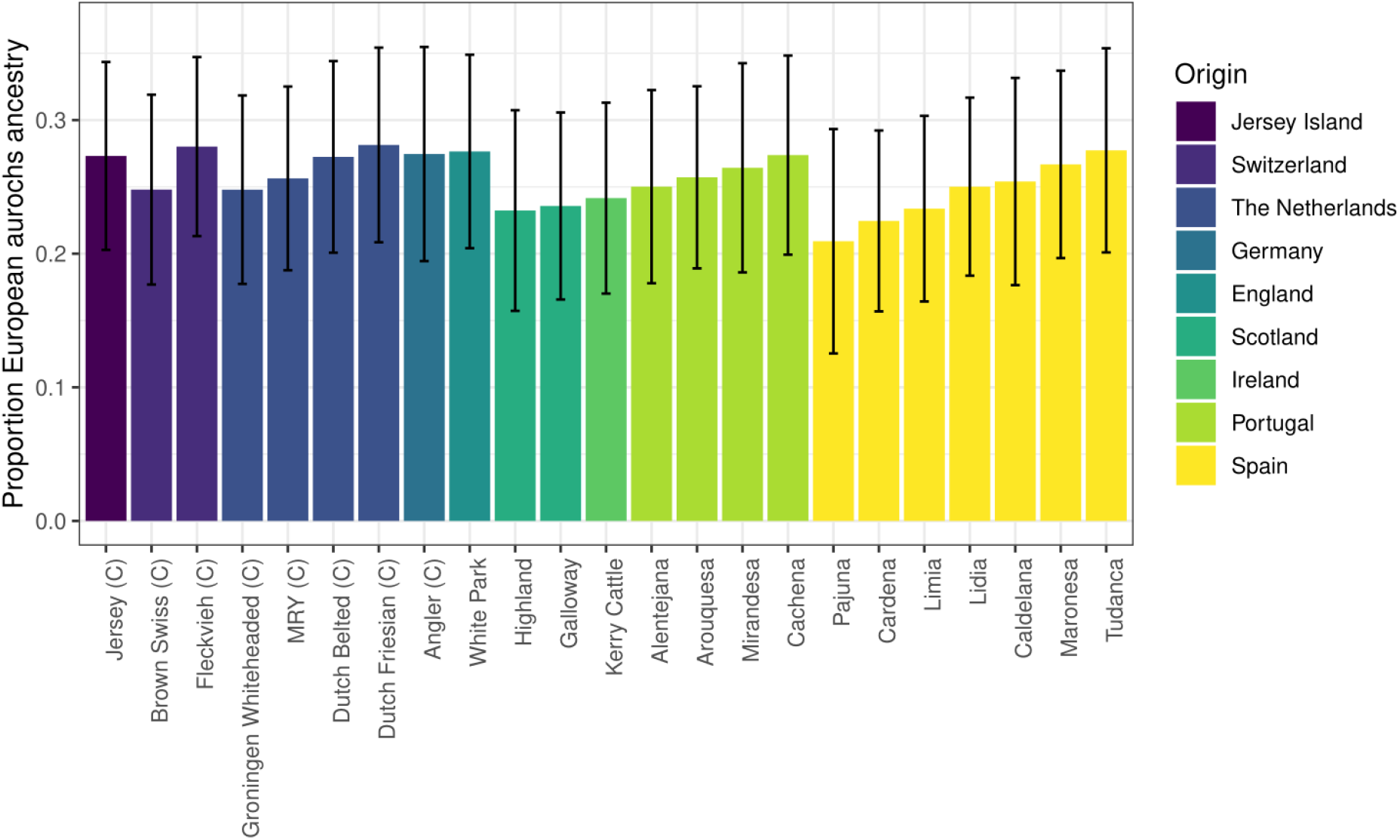
qpAdm estimates of Aurochs ancestry in modern western European cattle breeds from the (*27*) dataset. Commercial breeds are marked with a “C”. The figure is only showing breeds with feasible and non-rejected two source models, all results are shown in Supplementary Information, Table S1. Error bars are showing block-jackknife estimates of the 95% confidence interval.

Cattle have played an important role in Iberian culture during the last centuries as they have been part of numerous traditional popular events including bullfighting. The Lidia breed, a heterogeneous group of Iberian cattle that is mainly bred for aggressive behavior, has commonly been used for such popular festivities (*33*). Even though Lidia cattle has only been actively bred for agonistic behavior for about 200 years, some people attribute their aggressiveness and appearance as an indication of high levels of aurochs ancestry (*34*). We additionally use medium coverage genomes of six Lidia individuals (*41*) to estimate their proportion of aurochs ancestry. Lidia cattle in the (*27*) data set had a point estimate of 25% (95% confidence interval: [18.5, 31.5]) aurochs ancestry and estimates in the individual genomes ranged from 17.6% [10.9, 24.3] to 23.5% [17.0, 30.0] (Supplementary Information, Figure S8) – all overlapping with the observed range for other western European breeds. Despite some variation between individuals, which might be attributable to noise due to low coverage sequencing data in the reference populations, we do not observe a systematically elevated level of aurochs ancestry compared to other modern breeds or ancient samples since the Bronze Age. While these results reject the idea that the specifics of Lidia cattle can be attributed to a substantially increased genome-wide aurochs ancestry, it does not rule out the possibility that the roots of their aggressiveness and appearance are indeed due to aurochs variants at key loci responsible for those traits. An in-depth investigation of such questions would require a larger dataset of aurochs genomes as well as a more comprehensive Lidia sampling due to their fragmentation in highly distant genetic lineages (*33*).

### Stable Isotope Analysis

In addition to their ancestry, we studied the ecology of the bovids through stable isotope analysis of bone collagen. Lynch et al. (*48*) suggested that stable isotope data could be used to infer niche separation between the species in Britain, with domestic cattle in more open settings, while aurochs (about 1‰more depleted in *δ*^13^C) were habitually in more forested areas, or wet ground. This is most likely facilitated through human management of the domestic cattle, separating them from their wild counterparts. In contrast, Noe-Nygard et al. (*49*) failed to observe such an effect in samples from Denmark and northern Germany.

Considering our dataset and other data published on Iberian cattle (categorised on morphology/date) (*50–59*) we observe that the nitrogen isotope means are statistically different only when our data are compared using morphological characteristics, not genetic distinctions (see Dataset S1 and Supplementary Information). This difference is mostly due to some domestic cattle with *δ*^15^N values greater than 6.5‰. This could be explained by some taurine cattle having exclusive habitual access to high nitrogen isotope ratio resources. For example, human management such as corralling on manured ground, or feeding with manured crops, would produce this effect. Nevertheless, there is generally a large amount of overlap in the isotope values for the two groups suggesting that wild and domesticated groups often did not occupy different niches in Iberia.

## Discussion

We generated and analyzed biomolecular data from *B. primigenius* and *B. taurus* spanning more than 9000 years in the same region. Cattle are important livestock in the Iberian Peninsula today, and our results illustrate the interaction between domestic cattle and their wild relatives in the past. The two groups show signs of frequent hybridization starting soon after the arrival of cattle to the peninsula, as evident in our oldest directly dated Neolithic individual (moo039, 7426-7280 CalBP) where signals of carrying both ancestries are clear. Throughout the Neolithic, we observed large variations in the wild versus domestic ancestry per individual, but this pattern later stabilized (to 20-30% aurochs ancestry) from the Chalcolithic/Bronze Age onwards. As we do not know whether the sequenced individuals were hunted or herded, this could reflect a transition from hunting and herding to predominantly herding and it is possible that systematic herd management led to the nearly constant levels of aurochs ancestry over the last 4000 years. This period also coincides with several other societal changes; including the Bell Beaker complex and the introduction of human ancestry from the Pontic steppe into the Iberian Peninsula (*60–62*). Around this time, humans also started processing a significantly higher amount of dairy products connected with the “secondary product revolution” (*63, 64*). Aurochs were probably present in Iberia until Roman times (*31*) leaving possibilities for interbreeding but we cannot exclude that various factors such as hunting or changing vegetation had led to a substantial decline in the wild aurochs population around the early Bronze Age. A previous study on cattle morphology from the site of El Portalón described a decrease in size from the Neolithic to the Chalcolithic and a further significant size decrease from the Chalcolithic to the Bronze Age (*65*) and associated this change in size to the aridification of the area at this time (*66*). Indeed, this climatic change could also be related to a reduction of the aurochs population contributing to the stabilization of the levels of ancestry in domestic cattle from the Bronze Age to the present. Nonetheless, our stable isotope results suggest that wild and domesticated groups often did not occupy substantially different niches on the Iberian Peninsula. Material excavated from Denmark suggested that aurochs changed their niches over time (*49*) demonstrating some flexibility depending on local vegetation and the possibility of aurochs adapting to changing environments.

The reduced level of aurochs ancestry on the X chromosome (compared to the autosomes) in admixed individuals suggests that it was mostly aurochs males who contributed wild ancestry to domestic herds, a process that had been suggested based on the distribution of mitochondrial haplotypes before (*5*). A recent parallel study based using ancient genomes also detected male-biased aurochs introgression using similar methods as our study (*28*). Consequently, the offspring of wild bulls and domestic cows could be born into and integrated within managed herds. It is unclear how much of this process was intentional but the possibility of a wild bull inseminating a domestic cow without becoming part of the herd suggests that some level of incidental interbreeding was possible. For Neolithic Turkey, it has been suggested that allowing insemination of domesticated females by wild bulls was intentional, maybe even ritual (*67*). Modern breeders are still mostly exchanging bulls or sperm to improve their stock which manifests in a lower between-breed differentiation on the X chromosome (*46*).

The lack of correlation between genomic, stable isotope and morphological data highlights the difficulties of identifying and defining aurochs to the exclusion of domestic cattle. All of these data measure different aspects of an individual: their ancestry, ecology or appearance, respectively. While they can give some indication, none of them are a direct measurement of how these cattle were recognised by prehistoric humans or whether they were herded or hunted. It remains unclear whether our ancestry inferences had any correlation to how prehistoric herds were managed and how much intentional breeding is behind the observed pattern of hybridization. It is even possible that all hybrids identified in this study were part of domestic herds.

Even though wild aurochs populations went extinct, European aurochs ancestry survived into modern cattle with a relatively uniform distribution across western European breeds. Isolated Iberian Lidia, bred for their aggressiveness, appears to be no exception to this pattern. This rejects the notion that an overall increased proportion of aurochs ancestry causes the distinctiveness of certain breeds, but considering the functional relevance of archaic introgression into modern humans (*68*), it is possible that aurochs variants at functional loci may have a substantial influence on the characteristics of modern cattle breeds. Our low coverage sequencing data did not allow us to investigate this but future bioarchaeological studies combining different types of data will have the possibility to clarify the role of the extinct aurochs ancestry in modern domestic cattle.

## Conclusions

Using a bioarchaeological approach we have demonstrated that since cattle arrived in Iberia there has been hybridization with the local aurochs population, and that mainly aurochs bulls contributed to the gene pool still found in domestic herds today. Admixture proportions vary for the first few millennia but stabilize during the Bronze Age at approximately 20-30% of wild ancestry in the individuals found at the Iberian archaeological sites, a level that is still observed in modern Iberian breeds, including the more aggressive Lidia breed. This development could be the result of an initial mix of hunting and herding together with a generally loose management of herds, becoming more controlled over time in combination with a reduced importance of hunting wild aurochs.

The amount of hybridisation observed in the ancient cattle makes it difficult to genetically define what a domestic or wild *Bos* is, bringing into doubt the validity of such categorisations. Our interpretation is made more difficult by the overlap in morphological and metric data, creating further difficulties in species determination (especially in hybrids) and niche sharing as revealed by stable isotopes. To some extent, our interpretation is moot, as the salient matter is, how did prehistoric humans interact with cattle? What was their sense of wild and domestic and hybridisation? While we have recognised individual hybrids, to what extent these were part of domestic herds or intentionally bred and managed is uncertain.

Another source of uncertainty in our determinations is the limited knowledge about the genetic diversity in European aurochs. Further regional (and temporally longitudinal) aurochs genomes would aid future genomic studies defining the genetic variation in the European aurochs population.

## Materials and Methods

### Data generation

We attempted DNA extractions of 50 archaeological remains from which we successfully extracted DNA from 24 individuals identified as domestic cattle and aurochs excavated from four prehistoric sites in Iberia: El Portalón de Cueva Mayor (n=18), Artusia (n=1), Els Trocs (n=2) and Mendandia (n=3). Teeth and bones were UV irradiated (6 J/cm2 at 254 nm) and the first millimeter of bone/tooth surface abraded using a Dremel™ tool. DNA was extracted in a dedicated ancient DNA facility using a silica-based DNA extraction protocol (*69*). For each sample, 100-200mg of bone or tooth powder were incubated for 24 h at 37ºC, using the MinElute column Zymo extender assembly replaced by the High Pure Extender Assembly (Roche High Pure Viral Nucleic Acid Large Vol. Kit) and performed twice for each sample. DNA extracts were subjected to UDG treatment for the removal of deaminated cytosines and were further converted into blunt-end double stranded Illumina multiplex sequencing libraries (*70*). Between seven and fifthteen qPCR cycles were performed to amplify the DNA libraries using indexed primers (*70*). These were subsequently pooled at equimolar concentrations and shotgun sequenced on Illumina HiSeq and Novaseq sequencing platforms.

### Radiocarbon dates

Eight Bone and teeth were directly radiocarbon dated (AMS) at Waikato University in New Zealand and two teeth at Beta Analytics in the United States. Radiocarbon dates were calibrated using the OXcal 4.4 program (*71*) and the IntCal20 calibration curve (*72*). Three samples from the site of Mendandia were conventionally radiocarbon dated at Groningen (Netherlands) radiocarbon laboratory and calibrated as above.

### Stable Isotopes Analysis

Many of the samples analysed here were radiocarbon dated and stable isotope data (via IRMS) were generated in this process, to augment this data we also produced stable isotope data for some additional samples in this dataset, where they were available. The additional samples underwent bone collagen or tooth dentine collagen extraction at the Laboratorio de Evolución Humana (Universidad de Burgos) following the protocol of (*73*). In brief, this is a cold acid demineralization, followed by Milli Q water rinsing, gelatinization at pH3 (24 hrs at 70ºC), Ezee filtering and lyophilization. Collagen yields (as % mass of starting bone) were recorded. Stable isotope values (*δ*^13^C, *δ*^15^N) and %C, %N were measured in duplicate at the Universitat Autònoma de Barcelona, unless only one sample was successful in the analysis. Collagen samples (approx. 0.4 mg) were analysed using a Flash IRMS elemental analyser (EA) coupled to a Delta V Advantage isotope ratio mass spectrometer (IRMS), both from Thermo Scientific (Bremen, Germany) at the Institute of Environmental Science and Technology of the Universitat Autònoma de Barcelona (ICTA-UAB). International laboratory standard IAEA-600 was used, with measurements made relative to Vienna PeeDee Belemnite (V-PDB) for *δ*^13^C, and air N_2_ (AIR) for *δ*^15^N. The average analytical error was <0.2‰(1*σ*) as determined from the duplicate analyses of *δ*^13^C and *δ*^15^N. In house standards used was dog hair collected and homogenized for interlaboratory comparisons.

### Data processing

HiSeq X10 reads have been trimmed and merged using AdapterRemoval (*74*) while adapters for NovaSeq 6000 reads have been trimmed with cutadapt (*75*) and merging was performed with FLASH (*76*) requiring a minimum overlap of 11bp. Single-end reads of at least 35bp length were then mapped to the cattle reference genomes UMD3.1 (*77*) and Btau5 (*78*) using bwa (*79*) with the non-default parameters: -l 16500, -n 0.01, and -o 2. Different sequencing runs per sample were merged with samtools (*80*) and consensus sequences were called for duplicate sequences with identical start and end coordinates (*81*). Finally, reads with more than 10% mismatches to the reference genome were removed. Biological sex was assigned to the samples mapped to the Btau_5 reference genome (as UMD3.1 does not contain a Y chromosome assembly) using the R_x_ method (*82*) modified for 29 autosomes.

Mitochondrial contamination was estimated following the approach used by Green et al. (*83*) for hominins. We first identified nearly private mutations (less than 5% frequency in the 278 diverse mitogenomes used by MitoToolPy and dometree (*84*), obtained from Dryad https://doi.org/10.5061/dryad.cc5kn) in each individual and then used the proportion of non-consensus alleles at these sites to estimate contamination. We restricted this analysis to sites with at least 10x coverage, a minimum base quality of 30. Furthermore, transition sites with a C or G in the consensus mitogenome were excluded to avoid over-estimation due to post-mortem damage. Standard errors were estimated assuming a binomial distribution around the point estimate. Code used for this step can be found at https://github.com/GuntherLab/mt_contam_domestic_green

For comparative purposes, we also processed published data from (*5, 18, 19, 26*) using the same bioinformatic pipeline. Furthermore, we downloaded sequence data for six Spanish Lidia cattle (*41*), a single modern water buffalo (*Bubalus bubalis*, Jaffrabadi-0845) (*85*) and a single zebu cattle individual (Sha_3b) (*86*) and processed them with our ancient DNA mapping pipeline. To obtain a pseudohaploid Yak (*Bos grunniens*) sequence, we followed the approach by (*27*) splitting the Yak reference genome (*87*) contigs into 100bp fragments and mapping them to the UMD3.1 reference genome.

### Data Analysis

Mitochondrial consensus sequences were called using ANGSD (*88*) and the options -doFasta 2 -doCounts 1 -minQ 30 -minMapQ 30. Mitochondrial haplogroups were then assigned to the whole mitogenome sequences using the Python script MitoToolPy (*84*).

For population genomic analysis, we used a panel of SNPs derived from Run6 of the 1000 genomes project (*89, 90*). We obtained a list of SNPs from (*91*) and reduced the panel to biallelic SNPs of at least 10% minor allele frequency in the joint European *B. taurus*/Asian *B. indicus* dataset. Prior to genotype calling, all ancient BAM files were modified such that Ts in the first 5 bases of each fragment and As at the last 5 base pairs of each fragment have a base quality of 2. This approach allows to include more sites than excluding all transitions which are potentially affected by post-mortem damage. It produces highly correlated f4 statistics (Supplementary Information, Figure S2) and f_4_ ratios (Supplementary Information, Figure S3) but much lower standard errors in f_4_ ratios due to the larger total number of sites (Supplementary Information, Figure S3). To generate pseudohaploid representations of each individual, we randomly draw a single read with mapping and base quality of at least 30 at each SNP position. If the allele carried by the ancient individual was not one of the two known alleles, we removed the site from the panel. Using this approach, ∽9.1 million autosomal and 248K X chromosomal SNPs were genotyped in the ancient samples. To compare the ancient samples to a diverse set of modern cattle, we used the panel of modern European breeds presented by (*27*) which were genotyped at ∽770,000 SNPs. The ancient samples were genotyped the same way as for the 1000 Bulls project SNP panel.

To conduct an ordination of the nuclear data, sequences of 43 ancient Eurasian cattle and two aurochs were obtained from (*26*) and (*5*). Outgroup f_3_ statistics were calculated for all pairs of our Iberian *Bos* samples, using a Yak (*Bos grunniens*) genome as an outgroup, and a distance matrix for all samples was calculated as 1-f_3_. All f-statistics were calculated in R version 4.1.2 (*92*) package ‘admixtools2’ (*93*). The distance matrix was used to compute scores for non-metric multi-dimensional scaling (NMDS) ordinations using the metaMDS function in the ‘vegan’ R package and 10000 random starts (*94*).

European aurochs introgression ɑ into Iberian individual X was estimated using f_4_ ratios calculated with POPSTATS (*95*) and the equation

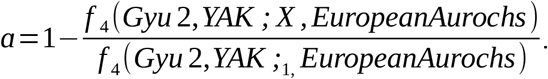

Both Bed3 and CPC98 were separately tested as aurochs source and Bed3 was chosen for the results presented in the article due to lower confidence intervals. POPSTATS was run with the non-default options –ratio, –testpop and –not23 to allow for more autosomes than humans have. We also used admixtools2 (*93*) and qpAdm (*38, 39*) to model the ancestry proportions in the samples. Bed3 was used as a source for European aurochs ancestry (due to lower standard errors in the f_4_ ratios) while the domestic Anatolian Neolithic Sub1 was used as a source for domesticated cattle ancestry. As “right” populations, we used Gyu2, *B. indicus*, Yak and *Bison bonasus bonasus* PLANTA. qpAdm was run with auto_only=FALSE, maxmiss=0.5 and allsnps=TRUE. When the two source model did not fit (p<0.01) or produced infeasible admixture proportions outside [0, 1], we used rotate_models and qpadm_multi to find alternative models adding CPC98 as an additional possible source or “right” population. qpAdm was also used for the modern western European breed panel from (*27*) adding Bes2 (*5*) to the “right” populations and excluding breeds from Italy and the Balkan from the targets as non-taurine ancestry (*27*) in them would lead to a rejection of the models. Finally, we also used Struct-f4 (*40*) to estimate ancestry proportions. First, input files were generated with the provided helper scripts and f_4_ statistics were calculated in blocks of 5Mbp. Struct-f4 was then run in semi-supervised mode to estimate ancestries in Iberian individuals with at least 0.1x coverage. This cutoff was chosen as lower coverage samples prevented conversion. CPC98, YAK, Ch22, Gyu2, Bed3, Sub1 and Sha_3b were used as additional individuals to provide a framework of different possible ancestries.

## Data availability

Raw sequence data and aligned reads for the new ancient individuals are available through the European Nucleotide Archive under accession number PRJEB63140. All metric and isotope data are available in Dataset S1.

## Acknowledgements

We are incredibly grateful to the excavation teams at the four archaeological sites. Sequencing was performed at The National Genomics Infrastructure (NGI) Stockholm. The computations and data handling were enabled by resources provided by the National Academic Infrastructure for Supercomputing in Sweden (NAISS) and the Swedish National Infrastructure for Computing (SNIC) at Uppmax, partially funded by the Swedish Research Council through grant agreements no. 2022-06725 and no. 2018-05973. This project was supported by grants from the Royal Physiographic Society of Lund (Nilsson-Ehle Endowments) to T.G. and C.V., Vetenskapsrådet (2017-05267) to T.G., Ramón y Cajal (RYC2018-025223-I) to C.V and a Beatriz Galindo Fellowship (BGS220-461AA-69201) to C.S.

