## Supplementary Information for "The Genomic Legacy of Aurochs hybridization in ancient and modern Iberian Cattle"

Torsten Günther[1,*], Jacob Chisausky[1], M. Ángeles Galindo-Pellicena[2] , Eneko Iriarte[3] , Oscar Cortes Gardyn[4], Paulina G. Eusebi[4], Rebeca García-González[3], Irene Urena[5], Marta Moreno[6], Alfonso Alday[7], Manuel Rojo[8], Amalia Pérez[3,] Cristina Tejedor Rodríguez[8], Iñigo García Martínez de Lagrán[9], Juan Luis Arsuaga[4], José-Miguel Carretero[3,10], Anders Götherström[5], Colin Smith[3,11,*], Cristina Valdiosera[3,11*]

1: Human Evolution, Department of Organismal Biology Uppsala University, Sweden
2: Centro Mixto UCM-ISCIII de Evolución y Comportamiento Humanos, Spain
3: Laboratorio de Evolución Humana, Universidad de Burgos, Spain
4: Universidad Complutense Madrid, Spain
5: Centre for Palaeogenetics, Stockholm, Sweden
6: Instituto de Historia - CSIC, Spain
7: Área de Prehistoria, University of the Basque Country, Spain
8: Department of Prehistory and Archaeology, Valladolid University, Spain
9: Departamento de Prehistoria y Arqueología, UNED, Spain
10: Unidad Asociada de I+D+i al CSIC Vidrio y Materiales del Patrimonio Cultural (VIMPAC).
11: Dept. of Archaeology and History, La Trobe University, Australia.

### Sample/sites description

#### Artusia

The Artusia rock shelter is located at Unzué, in the eastern part of Navarre (Spain). It is positioned at the southern slope of a narrow path excavated between the Alaiz Mountains on the north, and the Unzué Peak on the south, by the Artusia stream. Its geographical location is characterised by a natural communication route between the aforementioned mountains and the Ebro valley, and by its proximity to differentiated biotopes and landscapes, similar to other Mesolithic shelters in the Ebro valley.

The rock shelter is filled by a stratigraphic package cut lengthwise by the Artusia stream exposing the different archaeological levels. Consequently, the archaeological works carried out in the shelter was a rescue excavation to prevent the collapse of the stratigraphy. Five phases of occupation were defined during the excavations, all of them within the Mesolithic, represented by the same technological evolution defined in the Ebro valley for this period:

- Artusia I: 1st phase of the Mesolithic of Notches and Denticulates: 7461-7145 cal BC: Abundant faunal and flint remains were recovered together with charcoal remains derived from successive human occupations.
- Artusia II: 2nd phase of the Mesolithic of Notches and Denticulates: 6689-6507 cal BC: The stratigraphic units show recurring human occupations of seasonal character. The

presence of occupation floors and hearths and the materials recovered (large quantities of flint and faunal remains) indicate that these human occupations were very intense.

- Artusia III: 1st phase of the Geometric Mesolithic: 6598-6453 cal BC: In general, the characteristics of the occupation layers and the hearths are the same. However, in Artusia III the occupations seem to be less intense than in Artusia II.
- Artusia IV: 2nd phase of the Geometric Mesolithic. There are no diagnostic materials, just charcoal and fauna remains.
- Artusia V: 3rd phase of the Geometric Mesolithic: 6205-6009 cal BC: Artusia V presents occupational contexts and hearths similar to those of previous phases.

In addition, Artusia stands out for the paleoclimatic information of its stratigraphy. Two different climatic events are detected: the 8.5 ka BP event (ca. 6550-6450 cal BC) and the 8.2 ka BP climate event (ca. 6300-6140 cal BC). These events were cold and dry and had important consequences for the landscapes and for the human groups. In this context, Mesolithic communities most likely suffered a scarcity of prey in an increasingly open environment (aridity and deforestation), which potentially required hunting techniques to be adapted in order to suit this new environment, e.g., projectile points made using geometric microliths. Altogether, the archaeological and environmental analysis of the Artusia rockshelter reinforces the idea that these hunter-gatherer groups had to adapt their lithic technologies and their lifestyles and subsistence strategies to a changing environment during the last phase of the Mesolithic.

#### Els Trocs

Els Trocs Cave (San feliu de Veri/Bisaurri, Huesca) is located at an over 1,500 m. altitude in a high mountain environment in the Pyrenees of Huesca, in the region of Alta Ribagorza in Aragon. Specifically, it is located on the southern slope of a conical hill, very close to the town of San Feliu de Veri (Municipality of Bisaurri, Huesca), and next to a plateau called partida de la Selvaplana, to the north of the Turbón massif and close to the Axial Pyrenees, equidistant from the rivers Ésera and Isábena. The site location is a strategic point where transhumance paths (cabañeras) that have almost certainly been used since the Neolithic converge. The site stands out for its proximity to fresh high altitude pastures, as well as to two saline springs, one of them, La Muria, less than 1 km from the cave, rich in salts such as chlorine and sodium, essential for sheep feeding due to their absence from the herbaceous pasture.

Despite the difficult conditions of temperature and humidity of the cavity, a complex unaltered stratigraphy has been described documenting a sequence of human occupation throughout two millennia. Based on the stratigraphic sequence and a set of radiocarbon dates, up to five moments of occupation of the cave can be identified:

- Trocs I. This is the first phase of occupation of the site, dated to the beginning of the last third of the 6th millennium cal. BC. The bovine remains analyzed in this work belong to this cycle of use of the cave.
- Trocs II. Phase dated to the middle of the 5th millennium cal. BC characterized, above all, by the presence of important amortized hearths.
- Trocs III. Uniform stratigraphic horizon and of notable power that is formed along almost a millennium; between the first third of the 4th millennium, until the beginnings of the 3rd millennium cal. BC. The presence of numerous smaller households than the previous ones reinforces the recurrent occupations of very short duration.

- Trocs IV. Partial funerary deposit from the end of the Neolithic and beginning of the Chalcolithic (3rd millennium cal. BC).
- Trocs V. The last occupation of the cave corresponds to historical times and is the most superficial phase of the stratigraphy, where materials from the Imperial Roman period have been found, including remains of ceramics, glass, iron objects and two coins that would allow a dating in the Lower Empire (4th century AD).

The dating of the different occupational levels of the Els Trocs cave is supported by a series of radiocarbon dates that have been carried out for each phase of the site. These dates, based on samples or short-lived events (cereal seeds, domestic fauna or human bones), have allowed us to determine the chronology of the site (*1–4*).

#### El Portalón de Cueva Mayor

This archaeological site is located in the karstic system of the Sierra de Atapuerca in Burgos, northern Spain. Archaeological excavations at El Portalón cave have exposed a stratigraphic sequence comprising the Late Pleistocene and the Holocene. A detailed record of more than 90 radiocarbon dates for the entire stratigraphic sequence range from 30,000 BP to 1,000 BP (*5*–*7*). While the Late Pleistocene sediments show scarce archaeological activity, the Holocene period is characterized by intense human occupation from the Mesolithic to Medieval times (*7*). All chronocultural periods are represented by abundant archaeological artefacts such as stone tools, ceramics, worked bone etc. Human remains are present throughout the whole period, but domestic faunal remains are much more frequent. Ovicaprines, cattle and pigs are the three most abundant domestic species, respectively, for the Neolithic, Chalcolithic and Early Bronze Age, but this pattern changed by the Middle Bronze Age where cattle became more abundant than ovicaprines (*8*, *9*). Aurochs (*Bos primigenius*) remains have also been identified at the site all throughout the Neolithic, Chalcolithic and Bronze Age (*8*). The archeological evidence suggests that most of these domestic animals were used for human consumption (*9*), however, these have also been found as grave goods in a child burial during the Chalcolithic (*6*).

#### Mendandia

The prehistoric deposit of Mendandia was discovered in 1991 in the Upper Ebro River Basin, in the town of Sáseta (Treviño, Spain). It was discovered and excavated by A. Alday between 1992-1995 and in 1997: work was carried out on 13 square metres, which made it possible to differentiate a stratigraphy with five sedimentological levels and six cultural units. It is a continuous sequence, without erosive phases, whose characters derive from natural sedimentation and human activity, providing very dense archaeological collections (*10*). It is a rock shelter of medium dimensions (15 metres long by 5 metres wide) settled on a wide terrace of 27 metres by 14 metres. At its feet runs the river Ayuda, the great collector of the Treviño Basin. The catchment area is located on the Oquina-Sáseta Ravine, a natural north-south corridor, surrounded by valley, mid-mountain and even high altitude environments with peaks close to a thousand metres in altitude. The plant and animal resources available during prehistoric times were abundant and diverse, making it an ideal human settlement for several generations: two thousand years of uninterrupted visitation have been recorded.

Five lithostratigraphic horizons (from bottom to top V, IV, III, II and I) and six cultural stages have been described. The observations and different granulometric, mineralogical and chemical analyses carried out on the whole sequence and for well-chosen samples show a strong anthropic character to the fill (contributions of hunted fauna, tools, prepared fires).

The first occupations (Level V) date to 8500±60 BP (GrA-6874), and correspond to groups of small size and not very intense activities. Based on the characters and composition of the lithic instruments the complex is ascribed to the Mesolithic of laminar facies. The site was used for hunting horses, aurochs, goats, sheep, deer, roe deer and wild boar. Level IV is also Mesolithic but of the notched and denticulate facies with a chronology between 7810±50 (GrN-22744) and 7780±40 BP (GrN-22745). An unusually high number of 47,579 bone fragments were inventoried for this type of site: deer and roe deer were objects of special interest to the human group, but the importance of the aurochs is very striking, with 80% of them being less than two months old, which indicates a very specific hunting management strategy. The lithic industry includes 11,284 items, with 354 retouched objects: 35 scrapers, 58 perforators, more than a hundred and a half very characteristic notches and denticulates, 9 burins, among other retouched objects). The Mesolithic cycle ends with the last of the units, the geometric, present in Mendandia in level III-lower. Chronologically, the substitution of the campiñoid forms by the new microlithic ones is very fast moving into 7620±50 BP (GrN-22743). The Mesolithic cycle is replaced by the Neolithic, represented in Mendandia by levels III-superior, II and I. The antiquity of the horizons with dates of 7210±80 (GrN-19658) and 7180±45 BP (GrN-22742) (both for III-superior) and 6540±70 (GrN-22741) and 6440±40 BP (GrN-22740) for II and I respectively, is undoubtedly surprising. Although during the Neolithic the technological preferences vary, the use of the site does not change. The remains of fauna (horse, aurochs, goat, buck, deer, roe deer, wolf, fox, marten and rabbit) are still very numerous. There is no evidence of domestication. The occupation of the site ends after the early Neolithic with no further occupations.

Overall, almost 50,000 bone remains have been recovered, including horses, aurochs, goats, bucks, deer, roe deer and wild boar, as major species, but also wolves, foxes and martens. Based on the ages and sexes of the trapped animals, the hypothesis of a recurrent nomadism can be accepted: they would go to the refuge between late spring and early summer to harass mainly young and females, in a strategy with control of the hunted in order not to decimate the resources.

Hunting occupied much of the time of the inhabitants of Mendandia. The strategic location of the shelter enabled the groups to capture a wide variety of species: with certain variations in their percentages, 90% of the game hunted was roe deer, red deer and aurochs, followed, more distantly, by wild boar, goats, sheep, horses, foxes, wolves, badgers and wild cats, among others. The analysis of the age of the animals allows us to consider spring and early summer as the phase of greatest hunting activity. Moreover, the proportion of anatomical parts present and absent from each animal suggests that, in many cases, the meatier parts of the animals were consumed elsewhere by means of conservation practices such as smoking. The anthracological study would indicate the burning of wet wood for precisely this purpose.

### Data Analysis

#### DNA Data authentication

All samples included in the nuclear analysis show deamination patterns characteristic for authentic ancient DNA (Supplementary Figure S1).

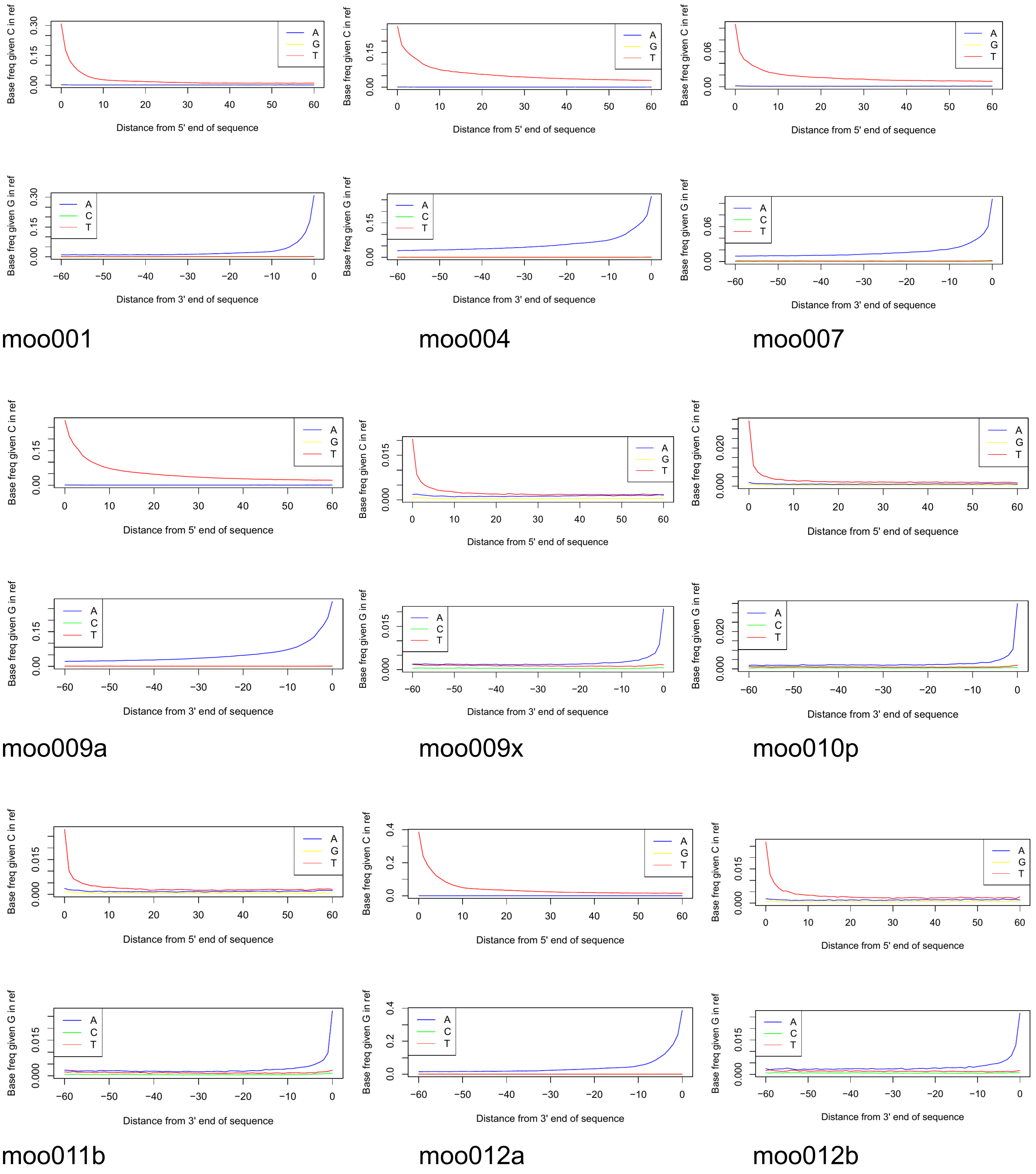

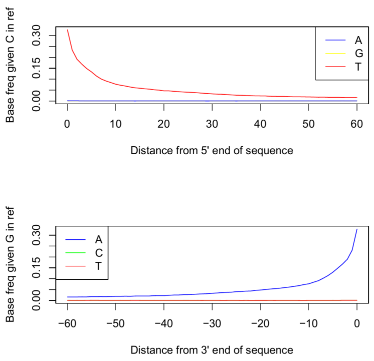

moo013a

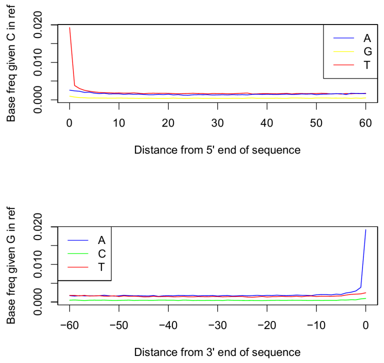

moo014

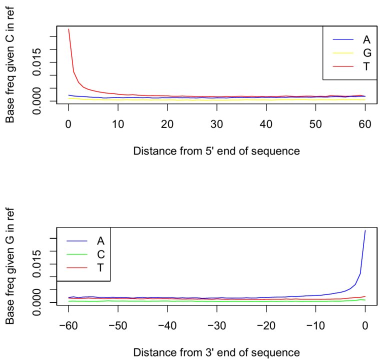

moo015

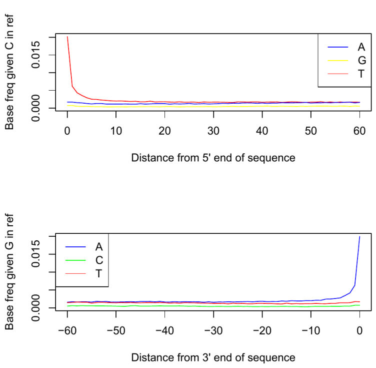

moo017

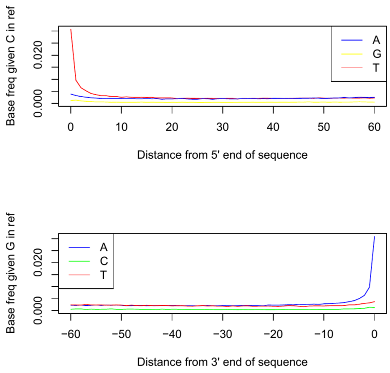

moo019

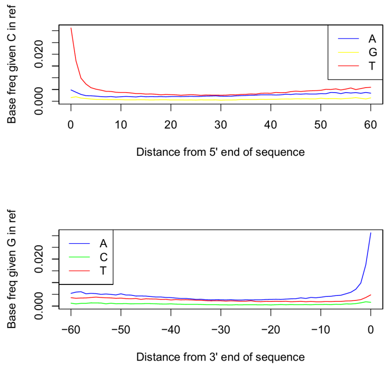

moo020

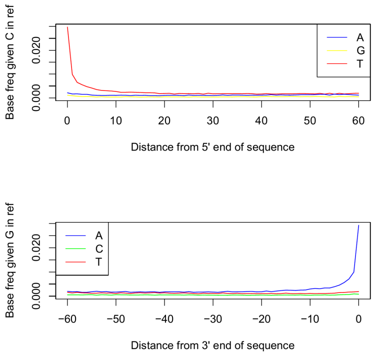

moo022

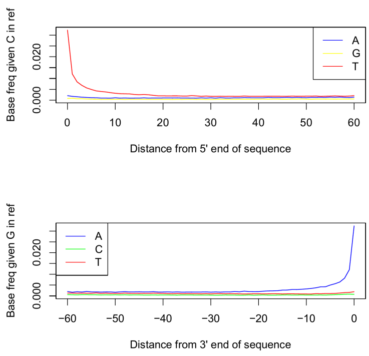

moo023

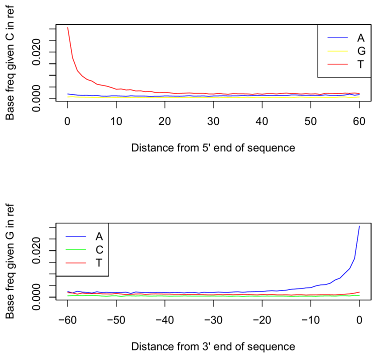

moo033

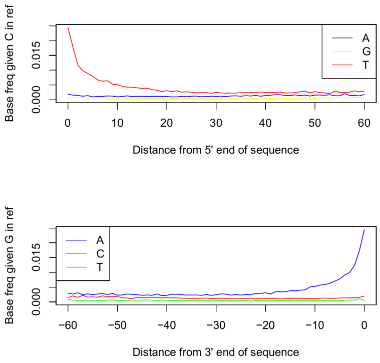

moo034

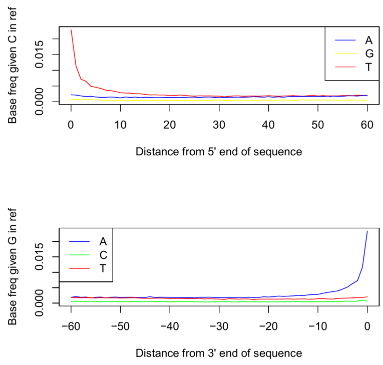

moo035

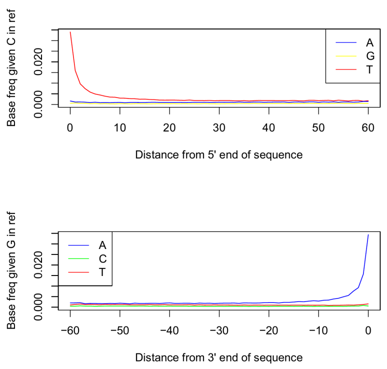

moo039

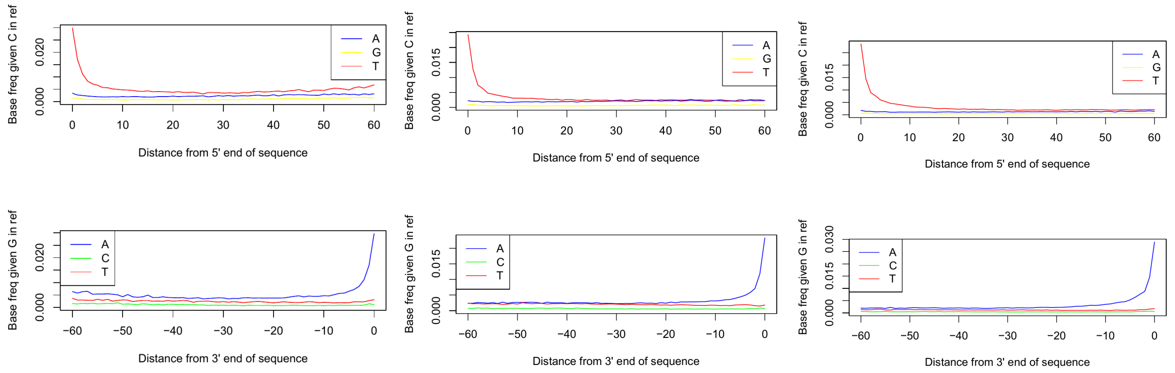

moo040 moo041 moo050

**Supplementary Figure S1**: Misincorporation plots for the samples included in nuclear analysis.

Prior to genotype calling, all ancient BAM files were modified such that Ts in the first 5 bases of each fragment and As at the last 5 base pairs of each fragment have a base quality of 2. As this is an approach that is not widely used, we compared its effect on downstream analysis. As our main analyses are all based on *f* statistics, we compared $f_4$ statistics (Supplementary Figure S2) and $f_4$ ratios (Supplementary Figure S3) of our rescaled base quality data with data only using transversion sites. While estimates are highly correlated, the data set reduced to transversions produces larger confidence intervals in $f_4$ ratios due to the lower number of sites (Supplementary Figure S3). Consequently, we decided to use the rescaled data for all analyses displayed in the main figures.

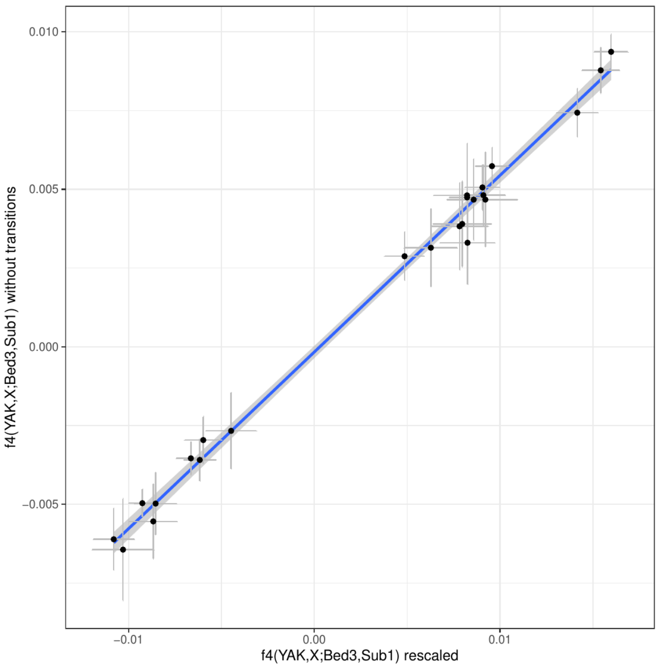

**Supplementary Figure S2**: $f_4$ statistics contrasting our newly sequenced samples to a reference domestic individual and a reference European aurochs. The $f_4$ statistics were calculated for two different versions of the bam files: with rescaled bases in fragment ends versus untreated bam files but a SNP panel excluding transitions. The blue line indicates a linear regression with confidence interval.

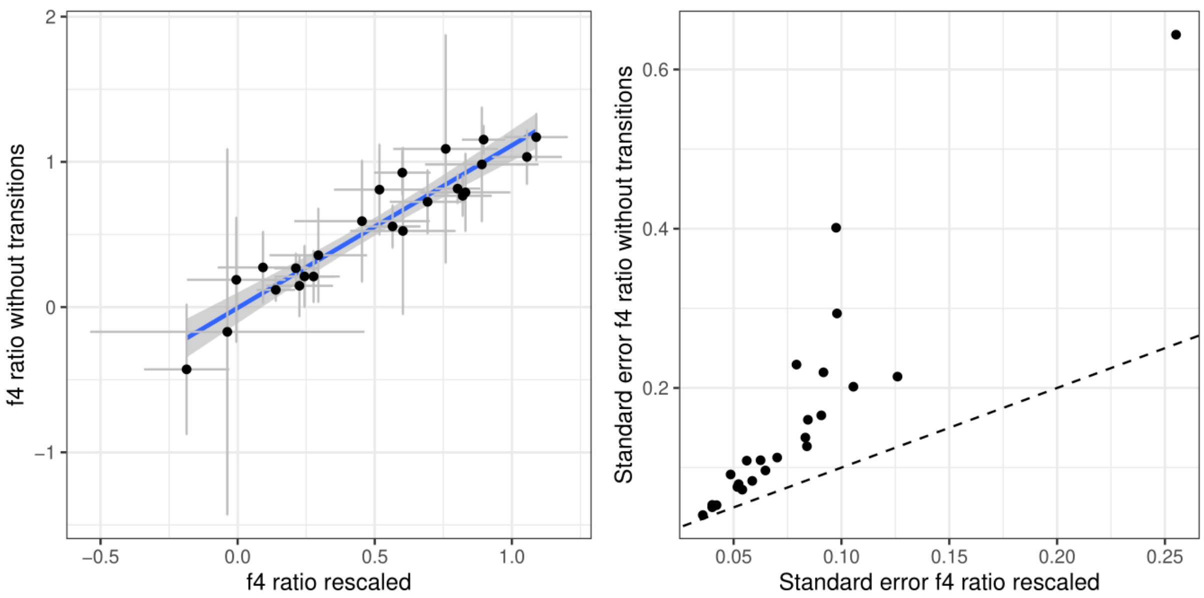

**Supplementary Figure S3**: $f_4$ ratios estimating European aurochs ancestry in our newly sequenced Iberian samples. The $f_4$ ratios were calculated for two different versions of the bam files: with rescaled bases in fragment ends versus untreated bam files but a SNP panel excluding transitions. The right panel shows the standard errors for the $f_4$ ratio estimates, the dashed line would correspond to equal standard errors for the two panels.

#### Principal Component Analysis

A principal component analysis (PCA) was conducted for the ancient samples with at least 0.01x coverage and modern breeds from (*11*). We used smartpca (*12*) with numoutlier: 0, killr2: YES, r2thresh: 0.4, lsqproject: YES and shrinkmode: YES to project the ancient samples onto the genetic variation defined by the modern breeds. Results are shown in Supplementary Figure S4 and S5. All Iberian samples fall close to the center of the plot, near modern central European breeds and the British aurochs CPC98. Some later samples appear to show stronger affinities to modern Iberian breeds. However, we do not see clear separation between predominantly aurochs or predominantly domestic individuals based on this analysis, highlighting the limitations of a PCA based on modern genetic variation if this is only a subset of the ancient variation that was present in the extinct wild ancestor.

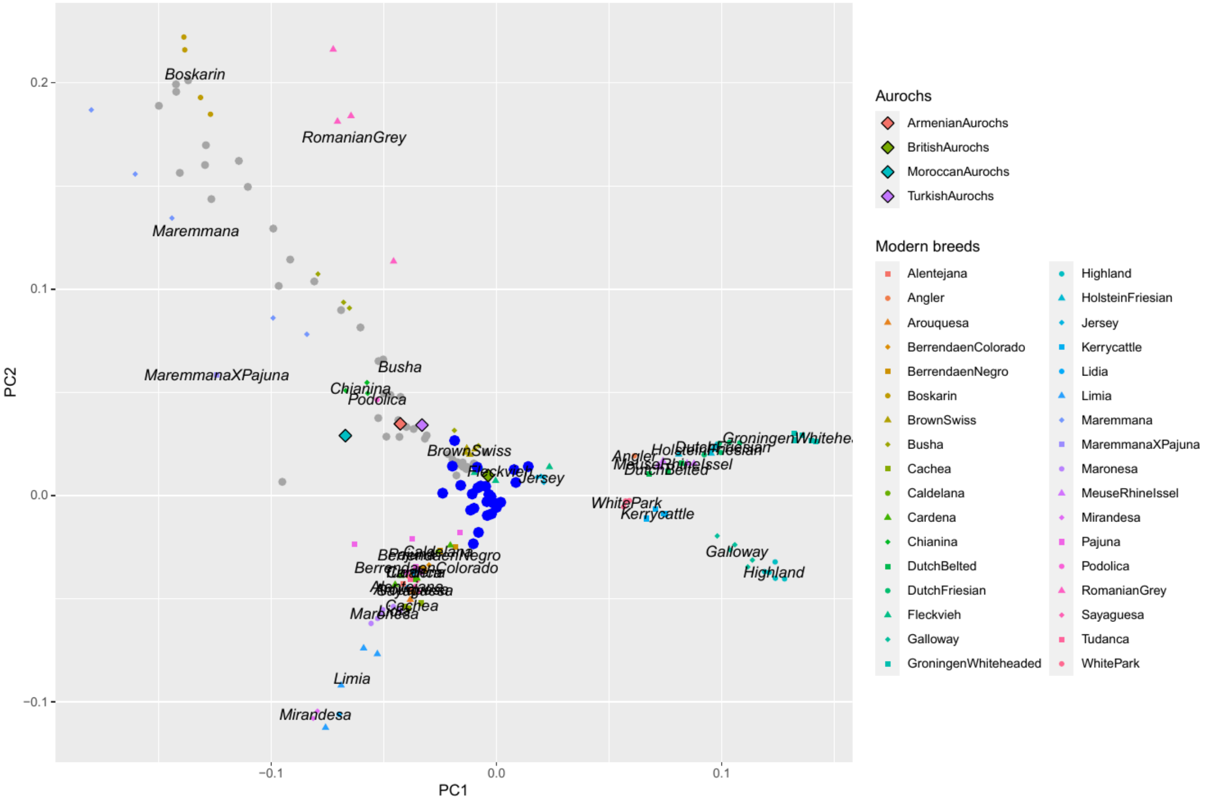

**Supplementary Figure S4** Biplot of PC1 and PC2 defined by modern western breeds together with ancient samples projected onto the PC-space. Aurochs genomes are shown as diamonds, ancient Iberian samples as blue dots and other ancient samples from (*13*) as grey dots.

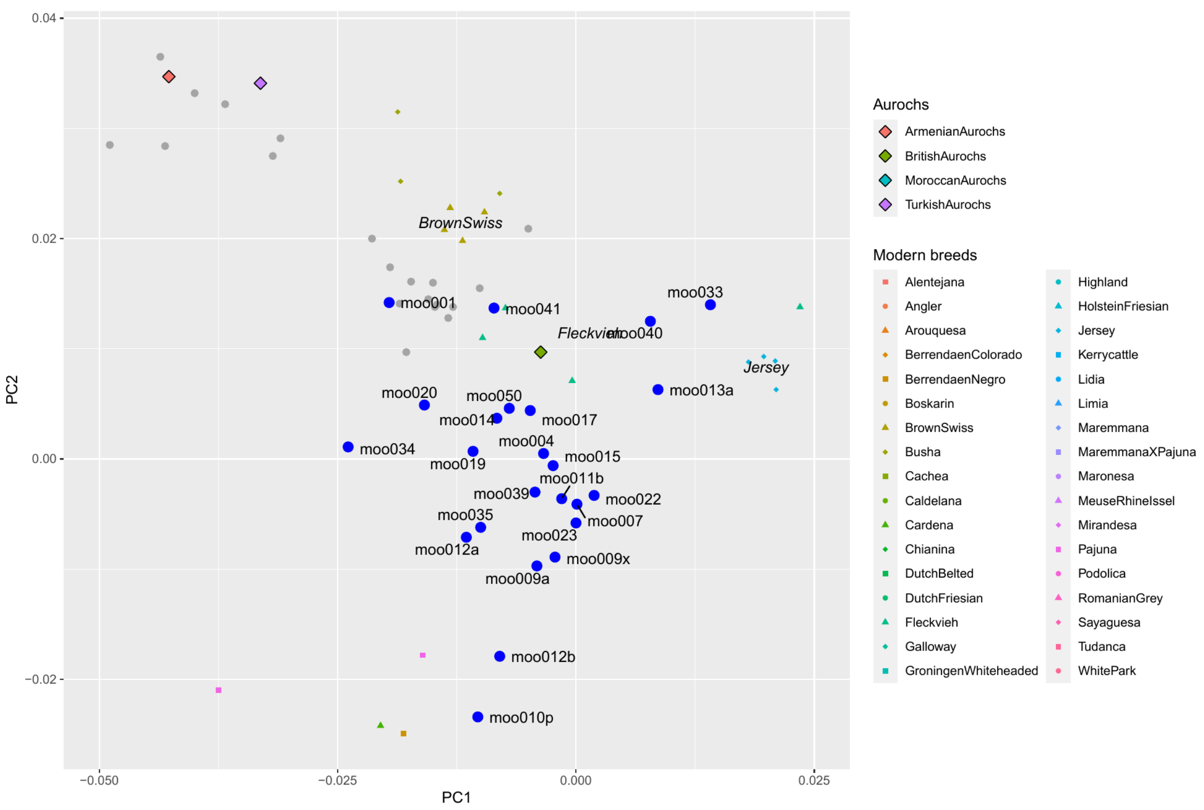

**Supplementary Figure S5**: Zoomed version of a biplot of PC1 and PC2 defined by modern western breeds together with ancient samples projected onto the PC-space. Aurochs genomes are shown as diamonds, ancient Iberian samples as blue dots and other ancient samples from (*13*) as grey dots. Sample IDs are added for Iberian samples.

#### Model based clustering

Unsupervised clustering of the ancient individuals together with modern breeds was carried out using ADMIXTURE (*14*, *15*). All individuals were haploidized by randomly drawing a single allele at each site to avoid artificial drift in the pseudohaploid ancient individuals. We then used plink (*16*, *17*) and the parameter --indep-pairwise 200 25 0.4 for linkage disequilibrium pruning. ADMIXTURE was run from K=2 to K=10 with 20 different seeds per K. Representative runs were then identified using pong in greedy mode with a similarity threshold of 0.95 (*18*). Results are displayed in Supplementary Figure S6. Similar to the PCA, the modern genetic variation dominates the cluster assignment not allowing for a differentiation between European aurochs, ancient European domestics and hybrid individuals. For higher numbers of clusters, there appear some differences in certain ancestry proportions between Iberian samples that are assumed to represent aurochs or domestic cattle (see other analysis) but all samples still carry ancestry from the same clusters and no cluster is exclusively representing European aurochs populations. This is likely a consequence of the dataset being heavily biased towards modern breeds which only represent a subset of the extinct aurochs variation.

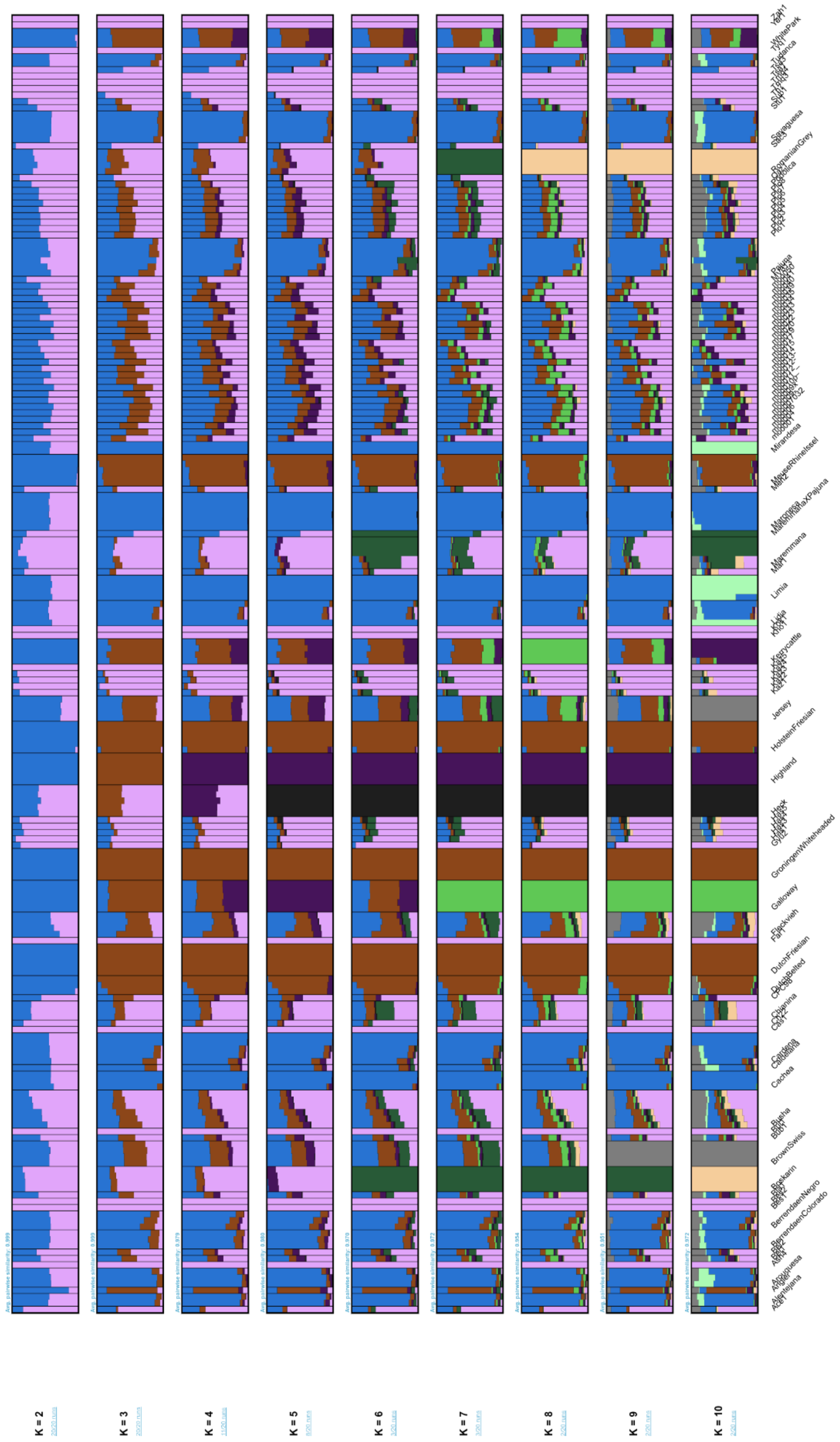

**Supplementary Figure S6**: Model-based clustering results for modern breeds and ancient Bos samples from K=2 to K=10.

#### Ancestry modelling in ancient Iberian samples

In addition to qpAdm and f4 ratios, we also used Struct-f4 (*19*) to estimate ancestry proportions in the ancient Iberian samples. We selected only ancient Iberian samples with at least 0.1x coverage and opted for a semi-supervised approach with the European aurochses Bed3 and CPC98, as well as YAK, Sha_3b (zebu cattle), Sub1 (Anatolian Neolithic), Gyu2 (Caucasus aurochs) and Ch22 (Anatolian aurochs) as reference samples.

At K=2, the two outgroups (Yak and indicine cattle) form a separate group (Supplementary Figure 7a), at K=3 each of them receive their own cluster (Supplementary Figure 7b) without any admixture in the Iberian samples. At K=4, the European aurochs (Bed3 and CPC98) separates from the domestic samples and western Asian aurochs. All tested Iberian samples now display non-zero proportions of both of these ancestries (Supplementary Figure 7c). At K=5, the Caucasian aurochs (Gyu2) splits into its own cluster with some Iberian samples displaying small proportions of this ancestry in addition to the two other components (Supplementary Figure 7d).

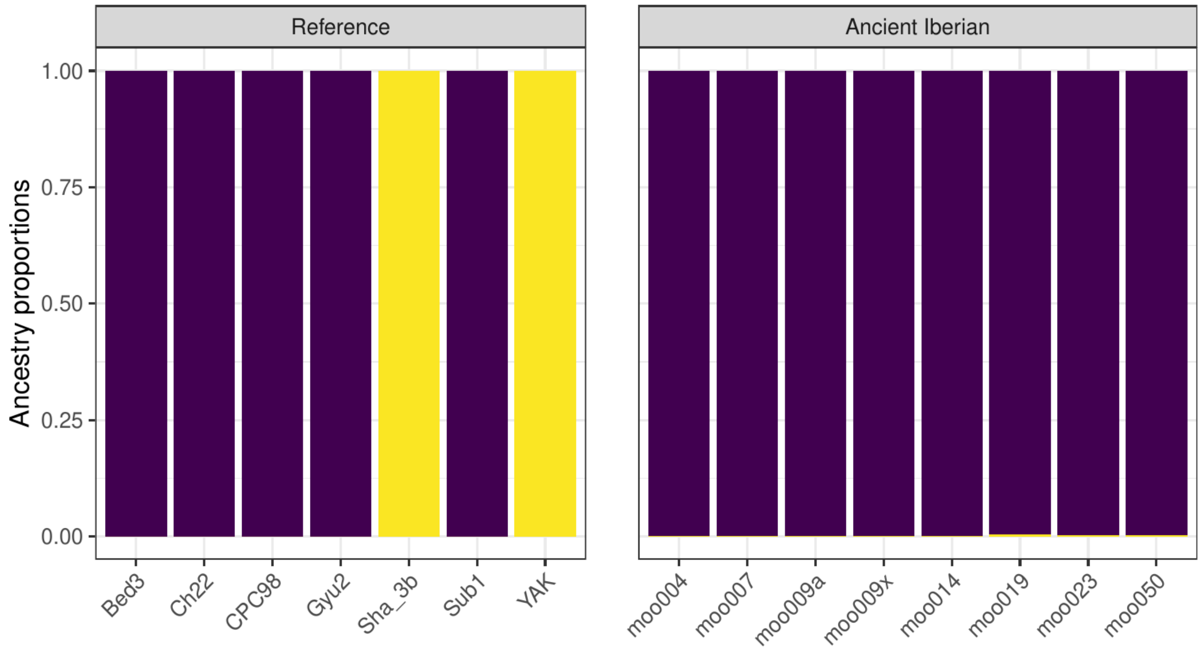

**Supplementary Figure 7a**: Structf4 results for K=2

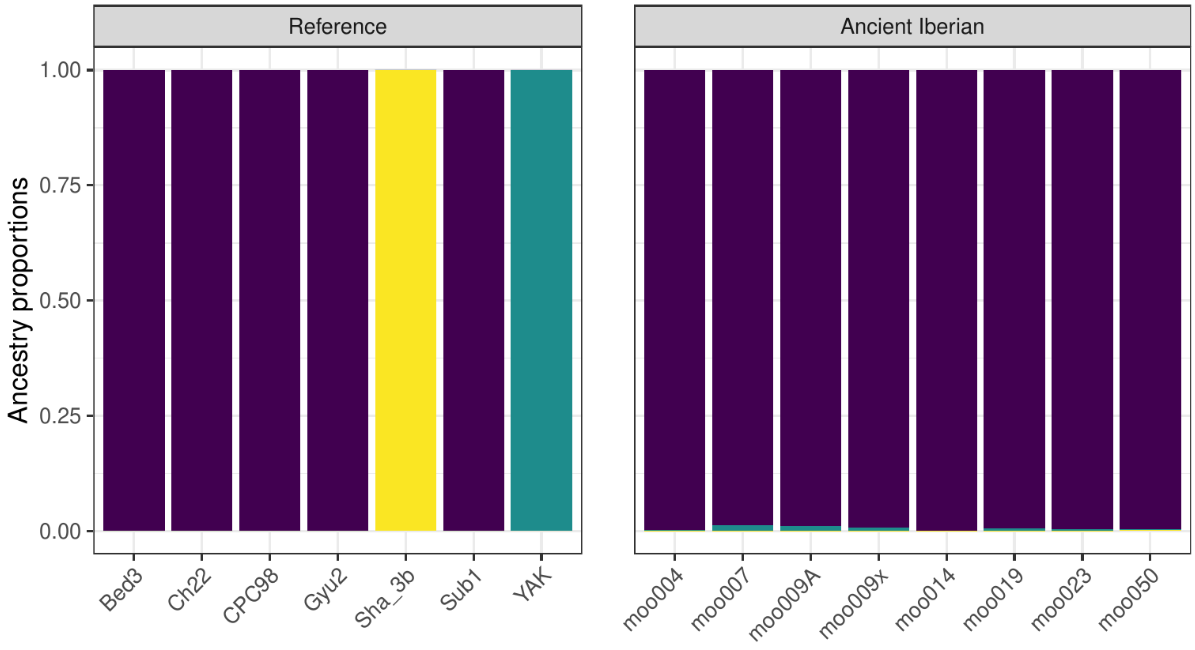

**Supplementary Figure 7b**: Structf4 results for K=3

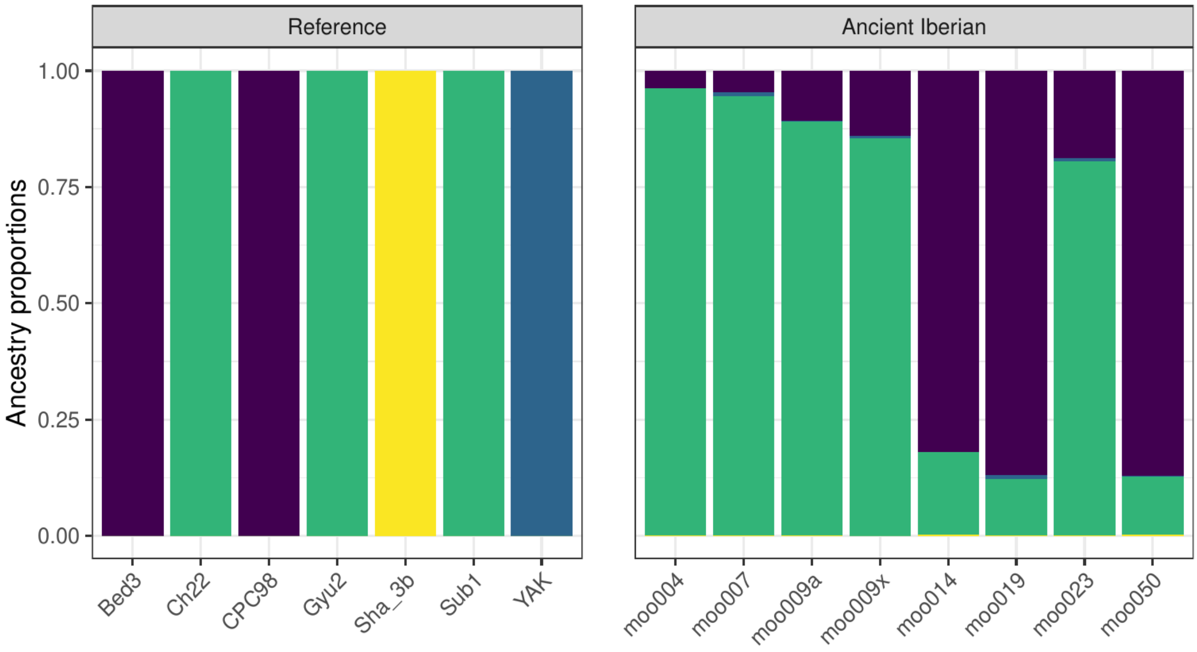

**Supplementary Figure 7c**: Structf4 results for K=4

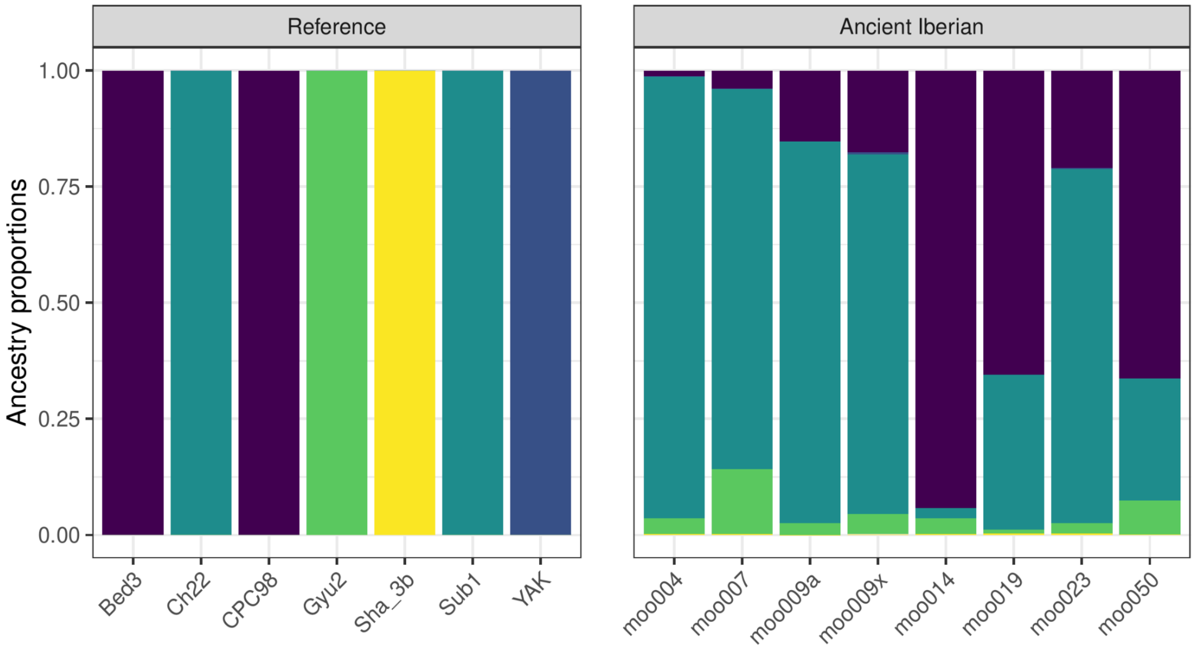

**Supplementary Figure 7d**: Structf4 results for K=5

#### Ancestry modelling in modern breeds

Since the publication of the first European aurochs genome, the Mesolithic British aurochs (CPC98) (*20*), several studies have contrasted different modern breeds in their amount of allele sharing with the aurochs genome (*11*, *20*, *21*) using *D* statistics (*22*). Such *D* statistics provide an assessment of relative allele sharing of two focal breeds with an outgroup but they do not directly estimate the amount of aurochs ancestry in each breed. The currently available data allows us to use more advanced methods that estimate the proportion of ancestries and are also able to reject certain ancestry models. Similar to the analysis of ancient samples, we used qpAdm (*23*, *24*) to model 30 modern European breeds with ancestry from two sources: Bed3 and Neolithic domestic Anatolian (Sub1). We only used western and central European breeds as target populations as southern and eastern European breeds showed signals of zebu cattle admixture (*11*) which would violate our two source model. Proportions of estimated aurochs ancestry are relatively similar across all breeds, ranging from 20.9% (Pajuna) to 28.1% (Dutch Friesian), see Supplementary Table S1. It is important to note that the standard errors of these estimates are all around 3%, so the 95% confidence intervals would be overlapping and the differences between breeds are not significant. The two source model was sufficient to explain the ancestry in 25 of the tested breeds ($p>0.01$) while the model was rejected for 5 breeds ($p<0.01$) including the commercial Holstein Friesian suggesting that they may contain small proportions of ancestries from outside of Europe. Estimates for the six individual Lidia genomes (*25*) are shown in Supplementary Figure S8.

**Supplementary Table S1**: qpAdm modelling of the genomic ancestry in modern European cattle breeds under a two source model using European aurochs (Bed3) and Anatolian Neolithic cattle (Sub1) as representatives of the sources. Genotypes for modern breeds from (*11*) (C = commercial breed)

| Breed code | Breed | Origin | Proportion European aurochs | Proportion Anatolia Neolithic | Standard error | P value |
|---|---|---|---|---|---|---|
| AL01 | Alentejana | Portugal | 0.2502 | 0.7498 | 0.0361 | 0.1030 |
| AN01 | Angler (C) | Germany | 0.2746 | 0.7254 | 0.0401 | 0.0747 |
| AR01 | Arouquesa | Portugal | 0.2572 | 0.7428 | 0.0341 | 0.0555 |
| BC01 | Berrenda en colorado | Spain | 0.2738 | 0.7262 | 0.0354 | *0.0077* |
| BN01 | Berrenda en negro | Spain | 0.3005 | 0.6995 | 0.0343 | *0.0014* |
| BS01 | Brown Swiss (C) | Switzerland | 0.2480 | 0.7520 | 0.0355 | 0.0343 |
| CA01 | Cardena | Spain | 0.2245 | 0.7755 | 0.0338 | 0.0141 |
| CC01 | Cachena | Portugal | 0.2738 | 0.7262 | 0.0372 | 0.0564 |
| CL01 | Caldelana | Spain | 0.2540 | 0.7460 | 0.0388 | 0.3139 |
| DB01 | Dutch Belted (C) | The Netherlands | 0.2724 | 0.7276 | 0.0358 | 0.2135 |
| DF01 | Dutch Friesian (C) | The Netherlands | 0.2814 | 0.7186 | 0.0364 | 0.0413 |
| FL01 | Fleckvieh (C) | Switzerland | 0.2801 | 0.7199 | 0.0335 | 0.0685 |
| GA01 | Galloway | Scotland | 0.2357 | 0.7643 | 0.0350 | 0.4203 |
| GW01 | Groningen Whiteheaded (C) | The Netherlands | 0.2479 | 0.7521 | 0.0353 | 0.3394 |
| HF01 | Holstein Friesian (C) | The Netherlands | 0.3402 | 0.6598 | 0.0352 | *0.0068* |
| HL01 | Highland | Scotland | 0.2323 | 0.7677 | 0.0375 | 0.0788 |
| JE01 | Jersey (C) | Jersey Island | 0.2732 | 0.7268 | 0.0352 | 0.3093 |
| KC01 | Kerry Cattle | Ireland | 0.2416 | 0.7584 | 0.0357 | 0.1596 |
| LI01 | Lidia | Spain | 0.2502 | 0.7498 | 0.0333 | 0.2204 |
| LM01 | Limia | Spain | 0.2337 | 0.7663 | 0.0347 | 0.3224 |
| ME01 | Maronesa | Spain | 0.2668 | 0.7332 | 0.0351 | 0.0339 |
| MI01 | Mirandesa | Portugal | 0.2643 | 0.7357 | 0.0391 | 0.0942 |
| MN01 | Maronesa | Spain | 0.2718 | 0.7282 | 0.0361 | 0.0400 |
| MR01 | MRY (C) | The Netherlands | 0.2564 | 0.7436 | 0.0344 | 0.0769 |
| PA01 | Pajuna | Spain | 0.2415 | 0.7585 | 0.0327 | *0.0097* |
| PA02 | Pajuna | Spain | 0.2093 | 0.7907 | 0.0420 | 0.0114 |
| SA01 | Sayaguesa | Spain | 0.2143 | 0.7857 | 0.0339 | *0.0021* |
| TU | Tudanca | Spain | 0.2773 | 0.7227 | 0.0382 | 0.1749 |
| TU01 | Tudanca | Spain | 0.2580 | 0.7420 | 0.0413 | 0.0295 |
| WP01 | White Park | England | 0.2765 | 0.7235 | 0.0362 | 0.5794 |

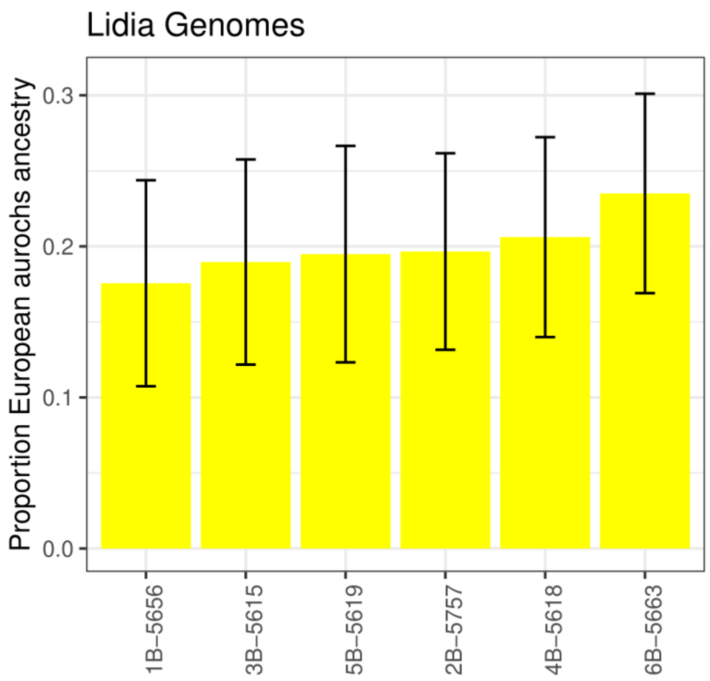

**Supplementary Figure S8**: qpAdm aurochs ancestry estimates for the six individual Lidia genomes.

We notice that these results differ from the D statistic results in the literature. (*11*) detected more aurochs alleles in breeds from the British Isles and Ireland, the Netherlands, Iberia and Jersey when compared to Alpine breeds (Brown Swiss and Fleckvieh) based on *D* statistics grouping the breeds by geographical origin. We can replicate these results on the breed level with most tests showing significantly more allele sharing between aurochs and the western European breeds relative to the Alpine breeds without any tests significantly pointing in the opposite direction (Supplementary Table S2). Furthermore, (*21*) presented *D* statistics suggesting more allele sharing between aurochs and Angus, Holstein and Jersey when compared to Iberian breeds. We can also largely reproduce this pattern by comparing Jersey and Holstein Friesian (Angus is not part of this study) with the Iberian breeds in our dataset (Supplementary Table S3): out of 28 individual tests, two short strong (Z>3) support and seven show intermediate support (Z>2) for more aurochs alleles in the non-Iberian breeds while none of the tests is showing significant support of the opposite pattern.

**Supplementary Table S2**: *D* statistics verifying the results of (*11*). pop3 is a Western European breed while pop4 is an Alpine breed. Negative values indicate an excess of allele sharing between pop2 and pop3.

| pop1 | pop2 | pop3 | pop4 | D | se | z | n |
|---|---|---|---|---|---|---|---|
| YAK | CPC98 | EL01 | FL01 | -0.02652393 | 0.00456532 | -5.80987815 | 450246 |
| YAK | CPC98 | EL01 | BS01 | -0.03070768 | 0.00487875 | -6.29417402 | 450241 |
| YAK | CPC98 | GA01 | FL01 | -0.03044061 | 0.00392968 | -7.74633205 | 450234 |
| YAK | CPC98 | GA01 | BS01 | -0.03442162 | 0.00434763 | -7.91732385 | 450223 |
| YAK | CPC98 | WP01 | FL01 | -0.02563453 | 0.00397421 | -6.45021264 | 450035 |
| YAK | CPC98 | WP01 | BS01 | -0.02957367 | 0.00450710 | -6.56157650 | 450031 |
| YAK | CPC98 | HL01 | FL01 | -0.02842438 | 0.00387999 | -7.32588339 | 450320 |
| YAK | CPC98 | HL01 | BS01 | -0.03246308 | 0.00432781 | -7.50104292 | 450319 |

| | | | | | | | |
|---|---|---|---|---|---|---|---|
| YAK | CPC98 | KC01 | FL01 | -0.02672545 | 0.00390891 | -6.83706658 | 450352 |
| YAK | CPC98 | KC01 | BS01 | -0.03070730 | 0.00431715 | -7.11286432 | 450340 |
| YAK | CPC98 | JE01 | FL01 | -0.01497177 | 0.00407495 | -3.67409596 | 450360 |
| YAK | CPC98 | JE01 | BS01 | -0.01907032 | 0.00441154 | -4.32282854 | 450342 |
| YAK | CPC98 | DB01 | FL01 | -0.00874258 | 0.00384590 | -2.27322063 | 450195 |
| YAK | CPC98 | DB01 | BS01 | -0.01281078 | 0.00438547 | -2.92118480 | 450193 |
| YAK | CPC98 | DF01 | FL01 | -0.01004034 | 0.00368917 | -2.72157043 | 450373 |
| YAK | CPC98 | DF01 | BS01 | -0.01412771 | 0.00421435 | -3.35228684 | 450371 |
| YAK | CPC98 | HF01 | FL01 | -0.00512796 | 0.00341739 | -1.50055117 | 450378 |
| YAK | CPC98 | HF01 | BS01 | -0.00914451 | 0.00385993 | -2.36908732 | 450376 |
| YAK | CPC98 | MR01 | FL01 | -0.01566191 | 0.00346923 | -4.51451546 | 450313 |
| YAK | CPC98 | MR01 | BS01 | -0.01967925 | 0.00393753 | -4.99786229 | 450307 |
| YAK | CPC98 | AL01 | FL01 | -0.00515036 | 0.00420798 | -1.22395152 | 450178 |
| YAK | CPC98 | AL01 | BS01 | -0.00916224 | 0.00476407 | -1.92319693 | 450170 |
| YAK | CPC98 | AR01 | FL01 | -0.00733482 | 0.00358002 | -2.04881890 | 450360 |
| YAK | CPC98 | AR01 | BS01 | -0.01140575 | 0.00397343 | -2.87050092 | 450357 |
| YAK | CPC98 | CC01 | FL01 | -0.01185512 | 0.00429875 | -2.75780703 | 450210 |
| YAK | CPC98 | CC01 | BS01 | -0.01592615 | 0.00467210 | -3.40877812 | 450196 |
| YAK | CPC98 | CL01 | FL01 | -0.00620692 | 0.00463521 | -1.33908070 | 449745 |
| YAK | CPC98 | CL01 | BS01 | -0.01051648 | 0.00500205 | -2.10243173 | 449745 |
| YAK | CPC98 | MI01 | FL01 | -0.00384595 | 0.00456479 | -0.84252575 | 450236 |
| YAK | CPC98 | MI01 | BS01 | -0.00801398 | 0.00496662 | -1.61356702 | 450221 |
| YAK | CPC98 | BC01 | FL01 | -0.00087993 | 0.00355382 | -0.24760203 | 450356 |
| YAK | CPC98 | BC01 | BS01 | -0.00498252 | 0.00395135 | -1.26096675 | 450345 |
| YAK | CPC98 | BN01 | FL01 | 0.00099552 | 0.00349974 | 0.28445469 | 450351 |
| YAK | CPC98 | BN01 | BS01 | -0.00310245 | 0.00405350 | -0.76537570 | 450348 |
| YAK | CPC98 | CA01 | FL01 | -0.00565510 | 0.00342051 | -1.65329057 | 450392 |
| YAK | CPC98 | CA01 | BS01 | -0.00977242 | 0.00389924 | -2.50623975 | 450386 |
| YAK | CPC98 | LI01 | FL01 | -0.00497786 | 0.00344966 | -1.44300022 | 450380 |
| YAK | CPC98 | LI01 | BS01 | -0.00909228 | 0.00392637 | -2.31569677 | 450374 |
| YAK | CPC98 | LM01 | FL01 | -0.01139534 | 0.00342571 | -3.32641246 | 450377 |
| YAK | CPC98 | LM01 | BS01 | -0.01553056 | 0.00399342 | -3.88903604 | 450373 |
| YAK | CPC98 | PA01 | FL01 | -0.00469758 | 0.00300994 | -1.56068935 | 450393 |
| YAK | CPC98 | PA01 | BS01 | -0.00883186 | 0.00363419 | -2.43021611 | 450384 |
| YAK | CPC98 | PA02 | FL01 | 0.00015340 | 0.00519442 | 0.02953081 | 449545 |
| YAK | CPC98 | PA02 | BS01 | -0.00393103 | 0.00545916 | -0.72008057 | 449548 |
| YAK | CPC98 | SA01 | FL01 | -0.00887584 | 0.00354305 | -2.50514043 | 450377 |
| YAK | CPC98 | SA01 | BS01 | -0.01299958 | 0.00401364 | -3.23885174 | 450369 |
| YAK | CPC98 | TU | FL01 | -0.00446867 | 0.00469083 | -0.95263975 | 449680 |
| YAK | CPC98 | TU | BS01 | -0.00849743 | 0.00509594 | -1.66748978 | 449674 |

**Supplementary Table S3**: *D* statistics verifying the results of (*21*). pop3 is an Iberia breed while pop4 is a central or north-western European breed. Positive values indicate an excess of allele sharing between pop2 and pop4.

| pop1 | pop2 | pop3 | pop4 | D | se | z | n |
|---|---|---|---|---|---|---|---|
| YAK | CPC98 | AL01 | HF01 | 0.00010984 | 0.00444032 | 0.02473730 | 450186 |
| YAK | CPC98 | AL01 | JE01 | 0.00975475 | 0.00476772 | 2.04599760 | 450158 |
| YAK | CPC98 | AR01 | HF01 | -0.00209568 | 0.00376061 | -0.55727039 | 450365 |
| YAK | CPC98 | AR01 | JE01 | 0.00757226 | 0.00436317 | 1.73549446 | 450336 |
| YAK | CPC98 | CC01 | HF01 | -0.00664020 | 0.00434678 | -1.52761461 | 450211 |
| YAK | CPC98 | CC01 | JE01 | 0.00298956 | 0.00493465 | 0.60583115 | 450192 |
| YAK | CPC98 | CL01 | HF01 | -0.00089510 | 0.00478336 | -0.18712725 | 449748 |
| YAK | CPC98 | CL01 | JE01 | 0.00890081 | 0.00511564 | 1.73992026 | 449721 |
| YAK | CPC98 | MI01 | HF01 | 0.00119432 | 0.00474148 | 0.25188685 | 450238 |
| YAK | CPC98 | MI01 | JE01 | 0.01099329 | 0.00508326 | 2.16264559 | 450206 |
| YAK | CPC98 | BC01 | HF01 | 0.00420509 | 0.00380025 | 1.10653083 | 450359 |
| YAK | CPC98 | BC01 | JE01 | 0.01401724 | 0.00423143 | 3.31264957 | 450329 |
| YAK | CPC98 | BN01 | HF01 | 0.00610700 | 0.00367932 | 1.65981831 | 450360 |
| YAK | CPC98 | BN01 | JE01 | 0.01585867 | 0.00427758 | 3.70739043 | 450327 |
| YAK | CPC98 | CA01 | HF01 | -0.00046112 | 0.00364117 | -0.12663996 | 450396 |
| YAK | CPC98 | CA01 | JE01 | 0.00926087 | 0.00415092 | 2.23104269 | 450370 |
| YAK | CPC98 | LI01 | HF01 | 0.00017668 | 0.00374294 | 0.04720225 | 450385 |
| YAK | CPC98 | LI01 | JE01 | 0.00994126 | 0.00422452 | 2.35322801 | 450356 |
| YAK | CPC98 | LM01 | HF01 | -0.00611705 | 0.00379450 | -1.61208547 | 450383 |
| YAK | CPC98 | LM01 | JE01 | 0.00356800 | 0.00415838 | 0.85802696 | 450352 |
| YAK | CPC98 | PA01 | HF01 | 0.00044748 | 0.00328790 | 0.13609781 | 450395 |
| YAK | CPC98 | PA01 | JE01 | 0.01022468 | 0.00374297 | 2.73170629 | 450366 |
| YAK | CPC98 | PA02 | HF01 | 0.00519730 | 0.00529190 | 0.98212286 | 449549 |
| YAK | CPC98 | PA02 | JE01 | 0.01507871 | 0.00565298 | 2.66739024 | 449529 |
| YAK | CPC98 | SA01 | HF01 | -0.00363095 | 0.00362183 | -1.00251585 | 450382 |
| YAK | CPC98 | SA01 | JE01 | 0.00607429 | 0.00435903 | 1.39349602 | 450359 |
| YAK | CPC98 | TU | HF01 | 0.00055343 | 0.00493685 | 0.11210270 | 449674 |
| YAK | CPC98 | TU | JE01 | 0.01044897 | 0.00524529 | 1.99206629 | 449653 |

These *D* statistic results are an apparent contradiction to the qpAdm results. However, some of these tests contain breeds for which the two source model was rejected by qpAdm (Supplementary Table S1) including a breed used as “reference” in the comparisons (Holstein Friesian). *D* statistics are known to be sensitive to certain biases (*26*) including ghost admixture, i.e. gene flow from an unsampled population. Considering the complex history of commercial cattle breeds, it is possible that these specific breeds have received parts of their ancestry from another source than just ancient domestic European cattle and European aurochs. We tested this by running another set of qpAdm models moving *B. indicus* from the “right” populations to the sources, i.e. testing each breed as a composition of Anatolian Neolithic cattle, European aurochs and zebu cattle (Supplementary Table S4). As a result, we now have fitting models for all breeds (p>0.01). The estimates of Zebu ancestry are all rather low <12%. We assume that these models do not necessarily measure zebu ancestry in all breeds but more generally detect non-European ancestry in commercial breeds. Estimates of aurochs ancestry vary much more than in the two source models which can be partly explained by the increased uncertainties (SEs up to 5%) which also has the consequence that all breeds still have overlapping 95% confidence intervals.

**Supplementary Table S4**: qpAdm results for the modern breeds in a three source model.

| Breed | European Aurochs proportion | European domestic proportion | indicus proportion | SE (aurochs) | SE (taurus) | SE (indicus) | p |
|---|---|---|---|---|---|---|---|
| AL01 | 0.1489 | 0.8332 | 0.0179 | 0.0477 | 0.0416 | 0.0086 | 0.1290 |
| AN01 | 0.1334 | 0.8403 | 0.0264 | 0.0505 | 0.0439 | 0.0096 | 0.6038 |
| AR01 | 0.1465 | 0.8333 | 0.0202 | 0.0427 | 0.0370 | 0.0081 | 0.2356 |
| BC01 | 0.1446 | 0.8303 | 0.0252 | 0.0433 | 0.0378 | 0.0080 | 0.5834 |
| BK01 | 0.0399 | 0.8771 | 0.0829 | 0.0489 | 0.0426 | 0.0093 | 0.0774 |
| BN01 | 0.1223 | 0.8447 | 0.0329 | 0.0430 | 0.0375 | 0.0080 | 0.1930 |
| BS01 | 0.0690 | 0.9010 | 0.0300 | 0.0477 | 0.0416 | 0.0088 | 0.1299 |
| BU01 | 0.0683 | 0.8212 | 0.1106 | 0.0465 | 0.0405 | 0.0088 | 0.1236 |
| BU02 | 0.0481 | 0.8906 | 0.0613 | 0.0464 | 0.0403 | 0.0086 | 0.1336 |
| CA01 | 0.1713 | 0.8153 | 0.0135 | 0.0426 | 0.0370 | 0.0079 | 0.1055 |
| CC01 | 0.1525 | 0.8241 | 0.0233 | 0.0492 | 0.0425 | 0.0092 | 0.5879 |
| CH01 | 0.0313 | 0.8667 | 0.1020 | 0.0492 | 0.0429 | 0.0092 | 0.1833 |
| CL01 | 0.1003 | 0.8756 | 0.0241 | 0.0518 | 0.0452 | 0.0094 | 0.2378 |
| DB01 | 0.1892 | 0.7951 | 0.0158 | 0.0449 | 0.0390 | 0.0084 | 0.4417 |
| DF01 | 0.1837 | 0.7962 | 0.0201 | 0.0444 | 0.0386 | 0.0084 | 0.4395 |
| FL01 | 0.1138 | 0.8590 | 0.0271 | 0.0427 | 0.0372 | 0.0080 | 0.1621 |
| GA01 | 0.2064 | 0.7859 | 0.0077 | 0.0432 | 0.0377 | 0.0080 | 0.6631 |
| GW01 | 0.1925 | 0.7969 | 0.0107 | 0.0442 | 0.0386 | 0.0081 | 0.3708 |
| HF01 | 0.1941 | 0.7799 | 0.0260 | 0.0430 | 0.0374 | 0.0080 | 0.0834 |
| HL01 | 0.1905 | 0.8023 | 0.0072 | 0.0459 | 0.0400 | 0.0087 | 0.0260 |
| JE01 | 0.1233 | 0.8527 | 0.0240 | 0.0458 | 0.0399 | 0.0086 | 0.3298 |
| KC01 | 0.2136 | 0.7775 | 0.0089 | 0.0438 | 0.0382 | 0.0081 | 0.5727 |
| LI01 | 0.1345 | 0.8454 | 0.0201 | 0.0433 | 0.0377 | 0.0081 | 0.6682 |
| LM01 | 0.1640 | 0.8229 | 0.0131 | 0.0429 | 0.0374 | 0.0081 | 0.5898 |
| MA01 | 0.0401 | 0.8589 | 0.1010 | 0.0441 | 0.0384 | 0.0087 | 0.1193 |
| ME01 | 0.1554 | 0.8229 | 0.0217 | 0.0432 | 0.0377 | 0.0081 | 0.3929 |
| MI01 | 0.1468 | 0.8312 | 0.0220 | 0.0489 | 0.0426 | 0.0090 | 0.6720 |
| MN01 | 0.1336 | 0.8423 | 0.0240 | 0.0458 | 0.0399 | 0.0086 | 0.1144 |
| MR01 | 0.1525 | 0.8276 | 0.0198 | 0.0443 | 0.0387 | 0.0082 | 0.3582 |
| PA01 | 0.1437 | 0.8364 | 0.0199 | 0.0412 | 0.0357 | 0.0078 | 0.1722 |
| PA02 | 0.1864 | 0.8000 | 0.0135 | 0.0513 | 0.0444 | 0.0098 | 0.6806 |
| PO01 | 0.0003 | 0.8877 | 0.1119 | 0.0557 | 0.0485 | 0.0112 | 0.6150 |
| RO01 | 0.0318 | 0.8844 | 0.0838 | 0.0461 | 0.0402 | 0.0090 | 0.3009 |
| SA01 | 0.1260 | 0.8532 | 0.0208 | 0.0430 | 0.0375 | 0.0079 | 0.1805 |
| TU | 0.1698 | 0.8105 | 0.0196 | 0.0500 | 0.0434 | 0.0094 | 0.3151 |
| TU01 | 0.1303 | 0.8449 | 0.0248 | 0.0532 | 0.0465 | 0.0096 | 0.2313 |
| WP01 | 0.1637 | 0.8184 | 0.0179 | 0.0461 | 0.0402 | 0.0085 | 0.5317 |

Consequently, we believe that the *D* statistic results are at least partly driven by low levels of non-European ancestry in many commercial breeds. qpAdm, in contrast, did not just allow us to obtain quantitative estimates of aurochs ancestry but also to identify cases where the two sources did not fit the data. The confidence intervals for each breed are still rather wide, highlighting the need for more high-quality reference data from European aurochs as well as from informative groups that can be used as "right" populations for such analyses.

#### Stable Isotope data

Previous studies comparing the stable isotope values of domestic cattle and aurochs have noted differences between the species. A study of material excavated from the UK (*27*) noted an overall difference of 1‰ in the carbon isotope values, with aurochs being the most depleted in carbon-13. There was overlap in the values for the two species, but at sites where both species were found this difference ranged from 0.3 to 1.8‰. The overlap in the nitrogen isotope values was large and difficult to interpret. (*27*) interpret their data as reflecting different niches for the two species with domestic cattle in more open settings, while the aurochs could be in more forested areas (the canopy effect on carbon stable isotopes), or in wet ground (that might have a similar effect of depleting carbon isotopes). Consequently, (*27*) were suggesting that stable isotope data could be used to infer niche separation between the species, based on human management of the domestic cattle, separating them from their wild counterparts. For material excavated from sites in Denmark and Northern Germany (*28*), the situation is less clear due to the overlap in the values. However, aurochs values change over time (-20‰ to -24‰), which might reflect environmental change. The distinctions between the domestic cattle and aurochs in the above studies were based on morphological and size differences.

By comparison, we have used the material here to explore niche separation and/or other differences between wild and domestic cattle in the Iberian context. The data produced from the stable isotope analysis is given in Dataset S1 - Stable isotopes, with additional previously published data. We have compared our data to that from published studies, spanning the Mesolithic to the Bronze Age (*29–39*). While the stable isotope data is directly comparable, it should be noted that these different studies have used different skeletal elements from each other (sometimes horn, tooth or different types of bone) depending on availability, and have relied on morphology/size to determine species and sometimes species was only determined as *Bos sp.* Thus, there will be some inconsistency between studies as to how species has been determined, ostensibly they may have used different criteria or subjective criteria. Dates for these samples were either given in the publications or have been assigned here based on dates from related/associated material (often human samples dated from the same sites). Approximate latitudes and longitudes have also been assigned to the sites. Samples with C:N ratios of 3.7 were excluded from further analysis and in addition, to constrain analysis to likely C3 ecosystems only, three *B. taurus* samples from southern Spain (S-EVA 9042, S-EVA 7377, S-EVA 7378) have been excluded as they likely consumed some $C_4$ plants in their diet (*35*, *36*). For this analysis, the samples from El Callado described in the publication as *Bos species* (*32*) have been considered as *B. primigenius*, as they date to the Mesolithic.

The samples in our analysis were also from varied skeletal elements and originally classified on morphology to *B. taurus, B. primigenius* or not classified. We can consider these as similar to those in the published literature, with "not classified" equivalent to *Bos sp*.

#### Analysis based on morphology

Comparing $\delta^{15}$N against $\delta^{13}$C, the data does not reveal any obvious distinction between the groups (see figure S9), other than the *B. primigenus* tend to have higher $\delta^{13}$C values and lower $\delta^{15}$N values than *B. taurus* (only one *B. primigenius* sample has a $\delta^{15}$N value above 6.4‰). Testing the normality of the distributions was carried out in R studio using Shapiro-Wilk and Anderson-Darling tests (using nortest package) (see table S5). In most cases, at least one data set to be used in statistical comparisons was considered non-normally distributed, so comparisons have been carried out using a Wilcoxon rank sum test (Mann-Whitney test). All statistical analysis here was performed in R studio, Version 2024.09.0+375.

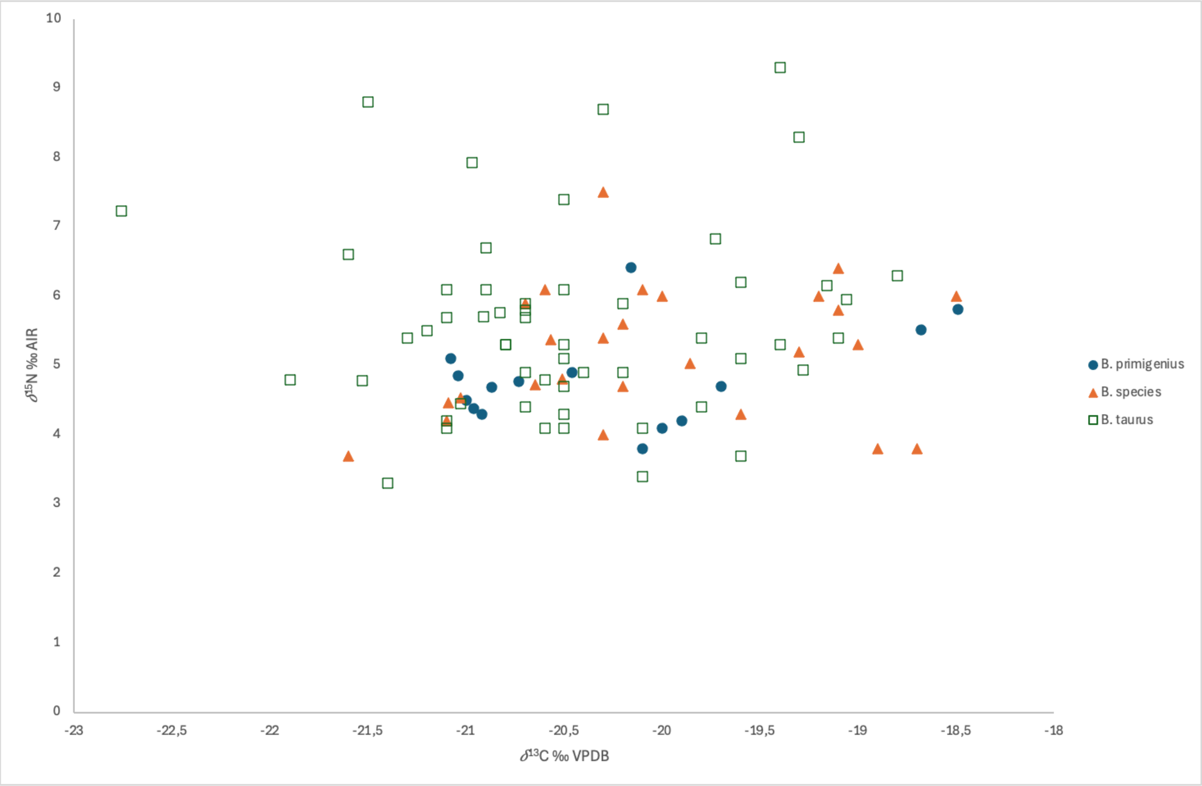

**Supplementary figure S9:** The distribution of $\delta^{15}$N vs $\delta^{13}$C for the samples, identified on the basis of morphology.

Wilcoxon rank sum tests of various groups, reveals no significant differences in the carbon isotope values between the groups tested (*B. primigenius* vs *B. taurus*, of when *B. primigenius* is included with likely *B. primigenius* and or tested against *B. taurus* and *B. species* as one group (see table S6 for W and p values). Statistically significant differences are observed when comparing nitrogen isotope values between (*B. primigenius vs B. taurus,*

*B. primigenius vs B. taurus* and *B. species* and when “likely *B. primigenius”* is added to the *B. primigenius* group in both comparisons). It should be noted that the stable isotope values are influenced by the environment and we can anticipate geographical differences in their distribution. Of importance here is that the geographic distribution of the species is not even across latitude and it can thus influence our observations. Six of the 13 aurochs (46%) are below 41º N, while only 3 of 54 (approx 6%) of *B. taurus* are below this latitude. The $\delta^{13}$C values are higher at lower latitudes and the distribution of carbon isotope values is narrower at lower latitudes as well, with no values lower than -20.1‰ observed south of 41º N. It should be noted most sites are north of 41º N and thus these differences could be a sampling artefact (especially the narrow range of values in the south), but as noted that the southern sites have some of the highest carbon isotope values and this would be expected given climatic/ecological differences between the north and south of Spain. Nevertheless, in the northern sites only (>41ºN), where most samples have been recovered the mean carbon stable isotope values of *B. primigenus* and *B. taurus* are not statistically different.

For nitrogen isotopes, there is no linear relationship between latitude and $\delta^{15}$N, and the range of values is similar across latitudes, except that values of 6.7‰ or greater are only observed north of 42º latitude. When comparing geographically only those animals recovered from >42ºN there are no statistical differences between groups for $\delta^{15}$N. The average $\delta^{15}$N value for all samples is 5.35‰ (sd 1.18), there are 11 *B. taurus* and *B. sp.* samples that fall outside the average + 1sd and no aurochs. This might reflect some level of management in the domestic fauna (corralling or feeding of manured crops or pastures).

**Supplementary Table S5:** Collagen isotope data summary based on morphological identification. Significant differences (at p=0.05) from normal distributions are highlighted in blue.

| | | | *$\delta$13C* | | | Normality tests | | | |
|---|---|---|---|---|---|---|---|---|---|
| | Morphological Species | Number of samples | Mean | Std Dev | Median | Shapiro Wilk W-value | Shapiro Wilk p-value | Anderson Darling A-value | Anderson Darling p-value |
| All | B. taurus | 54 | -20.48 | 0.80 | -20.55 | 0.972 | 0.242 | 0.628 | 0.097 |
| | B. species | 26 | -20.02 | 0.83 | -20.20 | 0.961 | 0.415 | 0.438 | 0.273 |
| | B. primigenius | 13 | -20.20 | 0.86 | -20.16 | 0.864 | 0.043 | 0.670 | 0.061 |
| | B. primigenius + B. primigenius? | 15 | -20.27 | 0.83 | -20.46 | 0.853 | 0.019 | 0.784 | 0.032 |
| | B. taurus + B. species | 80 | -20.33 | 0.83 | -20.50 | 0.976 | 0.133 | 0.810 | 0.035 |
| Northern (>41ºN) | B. taurus | 51 | -20.56 | 0.75 | -20.60 | 0.977 | 0.403 | 0.529 | 0.168 |
| | B. species | 21 | -20.26 | 0.73 | -20.30 | 0.955 | 0.415 | 0.429 | 0.282 |
| | B. primigenius | 7 | -20.82 | 0.31 | -20.92 | 0.775 | 0.023 | N/A | N/A |
| | B. primigenius + B. primigenius? | 9 | -20.80 | 0.31 | -20.92 | 0.835 | 0.050 | 0.662 | 0.055 |
| | B. taurus + B. species | 72 | -20.47 | 0.75 | -20.51 | 0.979 | 0.259 | 0.589 | 0.120 |

| | | | *$\delta$15N* | | | Normality tests | | | |
|---|---|---|---|---|---|---|---|---|---|
| | Morphological Species | Number of samples | Mean | Std Dev | Median | Shapiro Wilk W-value | Shapiro Wilk p-value | Anderson Darling A-value | Anderson Darling p-value |
| All | B. taurus | 54 | 5.58 | 1.33 | 5.40 | 0.940 | 0.010 | 0.963 | 0.014 |
| | B. species | 26 | 5.18 | 0.95 | 5.25 | 0.961 | 0.420 | 0.317 | 0.519 |
| | B. primigenius | 13 | 4.79 | 0.74 | 4.69 | 0.927 | 0.310 | 0.430 | 0.261 |

| | | | | | | | | |
|---|---|---|---|---|---|---|---|---|
| | B. primigenius + B. primigenius? | 15 | 4.80 | 0.69 | 4.70 | 0.938 | 0.361 | 0.397 | 0.324 |
| | B. taurus + B. species | 80 | 5.45 | 1.23 | 5.34 | 0.943 | 0.001 | 1.096 | 0.007 |
| Northern (>42ºN) | B. taurus | 46 | 5.60 | 1.41 | 5.40 | 0.935 | 0.013 | 0.975 | 0.013 |
| | B. species | 11 | 5.36 | 0.92 | 5.03 | 0.862 | 0.061 | 0.575 | 0.104 |
| | B. primigenius | 7 | 4.88 | 0.73 | 4.69 | 0.778 | 0.024 | N/A | N/A |
| | B. primigenius + B. primigenius? | 9 | 4.88 | 0.63 | 4.77 | 0.776 | 0.011 | 0.821 | 0.020 |
| | B. taurus + B. species | 57 | 5.55 | 1.33 | 5.38 | 0.930 | 0.003 | 1.291 | 0.002 |

**Supplementary Table S6:** Summary of Wilcoxon rank sum test results of comparisons of stable isotope data based on morphological characterisation. Significant diferences (at p=0.05) are highlighted in blue.

| | | | Wilcoxon rank sum test with continuity correction | |
|---|---|---|---|---|
| | Test comparison | | W-value | p-value |
| All | B. primigenius vs B. taurus | *δ13C* | 398 | 0.461 |
| | | *δ15N* | 213 | 0.029 |
| | B. primigenius vs B. taurus + B. species | *δ13C* | 549 | 0.752 |
| | | *δ15N* | 339 | 0.045 |
| | B. primigenius + B. primigenius? vs B. taurus | *δ13C* | 444 | 0.575 |
| | | *δ15N* | 246 | 0.021 |
| | B. primigenius + B. primigenius? vs B. taurus + B. species | *δ13C* | 607 | 0.947 |
| | | *δ15N* | 395 | 0.036 |
| Northern only | B. primigenius vs B. taurus | *δ13C* | 128 | 0.232 |
| | | *δ15N* | 105 | 0.141 |
| | B. primigenius vs B. taurus + B. species | *δ13C* | 165 | 0.135 |
| | | *δ15N* | 128 | 0.124 |
| | B. primigenius + B. primigenius? vs B. taurus | *δ13C* | 174 | 0.254 |
| | | *δ15N* | 136 | 0.106 |
| | B. primigenius + B. primigenius? vs B. taurus + B. species | *δ13C* | 223 | 0.131 |
| | | *δ15N* | 169 | 0.102 |

#### Analysis based on genetics

Considering the genetic analysis of our samples, some samples can be reclassified when compared with the morphological categories. Using the genetic data we have established 3 models for the categorisation of the animals for stable isotope data analysis. Model 1: categorises the animal on its majority genetic ancestry, so if an animal is greater than 50% aurochs, it is considered aurochs, otherwise domestic. Model 2: categorises the animal if it is in the top 30th percentile of its ancestry. So that 70% or above of aurochs ancestry = aurochs; 30% or below is considered taurine and anything in between is a hybrid. Model 3: is the same as 2 with a 20% cut-off. All other published data has been included in the analysis, retaining their morphological assignments to species. Our samples with too little DNA for analysis have been excluded. Making the same comparisons of the stable isotope data as above using the new categories, we can explore if the reassignment has made a difference (summarised in table S7). Similar to the morphological distinctions, no datasets pass both tests for data normality, so subsequent inferential analysis was made using Mann Whitney U tests (see table S8). Comparing *B. primigenius vs B. taurus,* defined using the models above, there are no statistical differences for any model considering all the data, nor only the northern datasets.

**Supplementary Table S7**: Collagen isotope data summary based on genetic (or morphological identification for published studies). Significant differences (at p=0.05) from normal distributions are highlighted in blue.

| | | | | δ13C | | | Normality tests | | | |
|---|---|---|---|---|---|---|---|---|---|---|
| | Genetic Model | Genetic Species | Number of samples | Mean | Std Dev | Median | Shapiro Wilk W-value | Shapiro Wilk p-value | Anderson Darling A-value | Anderson Darling p-value |
| All | Model 1 | B. taurus | 55 | -20.48 | 0.73 | -20.65 | 0.951 | 0.025 | 1.140 | 0.005 |
| | | B. primigenius | 17 | -20.38 | 0.99 | -20.46 | 0.937 | 0.283 | 0.470 | 0.216 |
| | Model 2 | B. taurus | 52 | -20.45 | 0.74 | -20.60 | 0.958 | 0.064 | 0.916 | 0.018 |
| | | B. primigenius | 16 | -20.23 | 0.81 | -20.31 | 0.884 | 0.045 | 0.612 | 0.092 |
| | | Hybrid | 4 | -21.41 | 0.90 | -21.03 | 0.719 | 0.019 | N/A | N/A |
| | Model 3 | B. taurus | 51 | -20.44 | 0.75 | -20.60 | 0.960 | 0.084 | 0.849 | 0.027 |
| | | B. primigenius | 11 | -20.08 | 0.89 | -20.10 | 0.893 | 0.150 | 0.478 | 0.187 |
| | | Hybrid | 10 | -20.93 | 0.76 | -20.94 | 0.826 | 0.030 | 0.867 | 0.016 |
| Northern (>41ºN) | Model 1 | B. taurus | 52 | -20.55 | 0.68 | -20.70 | 0.959 | 0.069 | 0.911 | 0.019 |
| | | B. primigenius | 11 | -20.87 | 0.76 | -20.92 | 0.852 | 0.045 | 0.74 | 0.038 |
| | Model 2 | B. taurus | 49 | -20.53 | 0.69 | -20.65 | 0.966 | 0.164 | 0.723 | 0.055 |
| | | B. primigenius | 10 | -20.68 | 0.45 | -20.90 | 0.851 | 0.060 | 0.636 | 0.068 |
| | | Hybrid | 4 | -21.41 | 0.90 | -21.03 | 0.719 | 0.019 | N/A | N/A |
| | Model 3 | B. taurus | 48 | -20.52 | 0.69 | -20.63 | 0.968 | 0.209 | 0.668 | 0.076 |
| | | B. primigenius | 5 | -20.81 | 0.37 | -20.92 | 0.763 | 0.039 | N/A | N/A |
| | | Hybrid | 10 | -20.93 | 0.76 | -20.94 | 0.826 | 0.030 | 0.867 | 0.016 |

| | | | | δ15N | | | Normality tests | | | |
|---|---|---|---|---|---|---|---|---|---|---|
| | Genetic Model | Genetic Species | Number of samples | Mean | Std Dev | Median | Shapiro Wilk W-value | Shapiro Wilk p-value | Anderson Darling A-value | Anderson Darling p-value |
| All | Model 1 | B. taurus | 55 | 5.48 | 1.31 | 5.30 | 0.915 | 0.001 | 1.424 | 0.001 |
| | | B. primigenius | 17 | 5.06 | 0.99 | 4.70 | 0.898 | 0.062 | 0.702 | 0.054 |
| | Model 2 | B. taurus | 52 | 5.51 | 1.33 | 5.30 | 0.918 | 0.002 | 1.325 | 0.002 |

| | | | | | | | | | | |
|---|---|---|---|---|---|---|---|---|---|---|
| | | B. primigenius | 16 | 4.93 | 0.85 | 4.70 | 0.914 | 0.133 | 0.557 | 0.126 |
| | | Hybrid | 4 | 5.50 | 1.31 | 5.15 | 0.877 | 0.326 | N/A | N/A |
| | Model 3 | B. taurus | 51 | 5.51 | 1.34 | 5.30 | 0.918 | 0.002 | 1.324 | 0.002 |
| | | B. primigenius | 11 | 4.77 | 0.81 | 4.50 | 0.893 | 0.151 | 0.572 | 0.106 |
| | | Hybrid | 10 | 5.40 | 0.99 | 5.10 | 0.885 | 0.148 | 0.480 | 0.179 |
| Northern (>42ºN) | Model 1 | B. taurus | 47 | 5.47 | 1.38 | 5.30 | 0.903 | 0.001 | 1.518 | 0.001 |
| | | B. primigenius | 11 | 5.26 | 1.05 | 4.90 | 0.816 | 0.015 | 0.897 | 0.014 |
| | Model 2 | B. taurus | 44 | 5.51 | 1.41 | 5.30 | 0.908 | 0.002 | 1.381 | 0.001 |
| | | B. primigenius | 10 | 5.07 | 0.87 | 4.80 | 0.799 | 0.014 | 0.875 | 0.015 |
| | | Hybrid | 4 | 5.50 | 1.31 | 5.15 | 0.877 | 0.326 | N/A | N/A |
| | Model 3 | B. taurus | 43 | 5.51 | 1.43 | 5.30 | 0.907 | 0.002 | 1.379 | 0.001 |
| | | B. primigenius | 5 | 4.87 | 0.88 | 4.50 | 0.700 | 0.010 | N/A | N/A |
| | | Hybrid | 10 | 5.40 | 0.99 | 5.10 | 0.885 | 0.148 | 0.480 | 0.179 |

**Supplementary Table S8**: Summary of Wilcoxon rank sum test results of comparisons of stable isotope data based on morphological characterisation (or morphological identification for published studies).

| | | | | Wilcoxon rank sum test with continuity correction | |
|---|---|---|---|---|---|
| | Test comparison | | | W-value | p-value |
| All | B. primigenius vs B. taurus | Model 1 | *δ13C* | 516 | 0.524 |
| | | | *δ15N* | 370 | 0.198 |
| | | Model 2 | *δ13C* | 478 | 0.374 |
| | | | *δ15N* | 298 | 0.089 |
| | | Model 3 | *δ13C* | 342 | 0.261 |
| | | | *δ15N* | 177 | 0.056 |
| Northern only | B. primigenius vs B. taurus | Model 1 | *δ13C* | 237 | 0.379 |
| | | | *δ15N* | 235 | 0.641 |
| | | Model 2 | *δ13C* | 217 | 0.578 |
| | | | *δ15N* | 178 | 0.349 |
| | | Model 3 | *δ13C* | 87 | 0.322 |
| | | | *δ15N* | 74 | 0.258 |

#### Conclusions

The values of the descriptive statistics of the stable isotope data here, are different depending on how our samples have been categorized and grouped with the published data, whether by morphology or genetics. However, using non parametric tests, a significant difference can only be observed when comparing between the $\delta^{15}$N values of *B. primigenius* vs *B. taurus* (and *B. primigenius vs B. taurus, B. primigenius vs B. taurus* and *Bos species* and when "likely *B. primigenius"* is added to the *B. primigenius* group in both comparisons) defined by morphology. When considering only sites north of 42ºN this difference is no longer significant. All other comparisons, whether using a morphological or a genetic characterisation of the individuals, are not significant.

The most obvious discriminating feature of the data is that of the distribution of $\delta^{15}$N values for taurine cattle and aurochs. This overlaps considerably, however, some taurine samples have $\delta^{15}$N values greater than 6.5‰, while aurochs samples do not. This can be interpreted as both types of cattle often share an ecological niche, but that some taurine cattle had habitual access to the areas or resources not available to the *B. primigenius*. This could indicate that in some instances the domestic cattle are managed by humans in certain ways (corralled on manured ground, fed manured crops?), but often not. It should be noted that there is wide variation in the $\delta^{15}$N within sites, so if this interpretation is true, not all cattle at a particular site are treated in the same manner. If we consider access to high nitrogen resources as a human management process, we can imply that aurochs are excluded from this, but that many cattle are in similar ecosystems as the aurochsen.

The observation here that the interpretation changes depending on the categorization of the samples represents a challenge to scientists using this kind of data. The genetic data can allow us to more easily observe what we can describe as aurochs-taurine hybrids and even quantify a degree of hybridisation, but ancient management strategies, may have been oblivious to the cattle ancestry, more concerned with the phenotypic attributes of the cattle (size temperament and manageability), in line with the morphological/metric and contextual assignment of species made in the present. Ancient farmers would have no knowledge of genetics, perhaps some knowledge of bloodlines, but would potentially herd/manage cattle that were useful and manageable, the latter might largely depend on size and temperament.

Contextualising our data with the UK and Danish/German datasets, there are some notable differences. In Northern Europe, average $\delta^{15}$N is lower in *B. taurus* relative to *B. primigenius* and the opposite pattern to that observed in Iberia. While the UK and Danish/German *B. taurus* have similar absolute $\delta^{15}$N values to those in Iberia. The Northern European individuals have lower $\delta^{13}$C values than their Iberian counterparts (attributable to climatic and ecological differences), and the *B. primigenius* in Northern Europe have the lower $\delta^{13}$C values compared to *B. taurus*. In Iberia this distinction between species is unclear. Considering only the northern Iberian sites *B. primigenius* is on average lower than *B. taurus* for $\delta^{13}$C, but considering all sites this is reversed. This may reflect similarities in the environment of Northern Iberia and the rest of Northern Europe (wetter and cooler) when compared to the south of Iberia. The significance of these differences is not clear, but considering the data as gross categories, it does appear that aurochs were confined to high nitrogen isotope areas in Northern Europe, but low $\delta^{15}$N areas in Spain, giving clear evidence for the need to interpret and contextualise such data regionally (this would need some reassessment, if in the future we find, evidence for long-distance movement of either species).
